# Evolutionary rescue and drug resistance on multicopy plasmids

**DOI:** 10.1101/2019.12.23.887315

**Authors:** Mario Santer, Hildegard Uecker

## Abstract

Bacteria often carry “extra DNA” in form of plasmids in addition to their chromosome. Many plasmids have a copy number greater than one such that the genes encoded on these plasmids are present in multiple copies per cell. This has evolutionary consequences by increasing the mutational target size, by prompting the (transitory) co-occurrence of mutant and wild-type alleles within the same cell, and by allowing for gene dosage effects. We develop and analyze a mathematical model for bacterial adaptation to harsh environmental change if adaptation is driven by beneficial alleles on multicopy plasmids. Successful adaptation depends on the availability of advantageous alleles and on their establishment probability. The establishment process involves the segregation of mutant and wild-type plasmids to the two daughter cells, allowing for the emergence of mutant-homozygous cells over the course of several generations. To model this process, we use the theory of multi-type branching processes, where a type is defined by the genetic composition of the cell. Both factors – the number of adaptive alleles and their establishment probability – depend on the plasmid copy number, and they often do so antagonistically. We find that in the interplay of various effects, a lower or higher copy number may maximize the probability of evolutionary rescue. The decisive factor is the dominance relationship between mutant and wild-type plasmids and potential gene dosage effects. Results from a simple model of antibiotic degradation indicate that the optimal plasmid copy number may depend on the specific environment encountered by the population.

## Introduction

Plasmids are extrachromosomal DNA elements that can be transmitted vertically or be transferred horizontally between cells and are commonly found in bacteria. Plasmids usually do not carry essential genes and are sometimes depicted as parasites that infect and exploit the bacterial cell, imposing a cost on its host. However, genes on plasmids can code for traits that are beneficial in specific environments such as resistance to antibiotics or the ability to metabolize rarely encountered carbon sources (Eberhard, 1989, 1990; MacLean and San Millan, 2015).

While some (usually large) plasmids only exist in a single copy within the bacterial cell, many plasmids are present in several copies per cell, some even in several hundreds (Friehs, 2004). The importance of the plasmid copy number for the evolutionary dynamics of genes on plasmids and the consequences for bacterial adaptation have recently started to gain increased attention (San Millan *et al*., 2016; Rodriguez-Beltran *et al*., 2018; Ilhan *et al*., 2019).

Compared to genes on a single-copy plasmid or on a haploid chromosome, there are several important differences. First, the mutational target size is multiplied by the copy number, making the appearance of mutations more likely (San Millan *et al*., 2016). Second, novel alleles appearing through mutation or novel genes acquired through transformation are initially only present on one of the plasmid copies. At cell division, as sketched in Fig. 1A, the segregation of plasmids to the daughter cells generates cells with a plasmid composition different from that of the mother cell, and thereby cells with a higher fraction of mutant plasmids may arise (San Millan *et al*., 2016; Halleran *et al*., 2019). Yet, through this segregation process, alleles on plasmids are subject to an extra-layer of drift, termed ‘segregational drift’ by Ilhan *et al*. (2019). This affects, in particular, the establishment probability of novel alleles on plasmids. On the other hand, the coexistence of mutant and wild-type plasmids within the same cell may allow the cell to escape from fitness trade-offs (Rodriguez-Beltran *et al*., 2018). Last, gene dosage effects can lead to an amplification of a gene’s effect such as increased levels of resistance if carried on a multicopy plasmid (Martinez and Baquero, 2000; San Millan *et al*., 2016; Santos-Lopez *et al*., 2017).

These effects have been demonstrated in several evolution experiments. San Millan *et al*. (2016) exposed *E. coli* populations that carried a bla_TEM-1_ *β*-lactamase gene either on the chromosome or on a multicopy plasmid to increasing levels of ceftazidime. Different alleles of the bla_TEM-1_ β-lactamase gene confer resistance to different antibiotics, and the original allele inserted by San Millan *et al*. (2016) conferred resistance to ampicillin but not to ceftazidime. Mutations can, however, generate an allele that provides ceftazidime resistance (see also Blazquez *et al*., 1995). San Millan *et al*. (2016) found that high levels of resistance were more likely to evolve when the gene was present on a multicopy plasmid due to (1) the increased mutational input, (2) the following increase of the fraction of mutant plasmids per cell through segregation, and (3) gene dosage effects. Using the same experimental system, Rodriguez-Beltran *et al*. (2018) showed that the existence of heterozygous cells, carrying wild-type and mutant plasmids, allowed populations to escape from fitness trade-offs, since these cells were resistant to both antibiotics. In a different setup, in the study by Santos-Lopez *et al*. (2017), the coexistence of a mutated plasmid and a wild-type plasmid allowed for a higher copy number and higher resistance than found for either plasmid type on its own, combining again two benefits of multicopy plasmids (the possibility of heterozygosity and gene dosage effects). Ilhan *et al*. (2019) finally studied the rate of evolution on two multicopy plasmids, one with a low and the other one with a high copy number. The accumulation of new mutations was lower than expected by the mutational target size, reflecting the decreased establishment probability of mutations on multicopy plasmids (at least in the absence of dosage effects).

Theoretical studies that quantify and disentangle the described effects are still rare. Some experiments are complemented by models and computer simulations (Rodriguez-Beltran *et al*., 2018; Ilhan *et al*., 2019). In addition, Halleran *et al*. (2019) show that random segregation of plasmid variants into the daughter cells speeds up adaptation compared to an equal partition that leads to exact copies of the mother cell.

In this article, we develop a mathematical framework to study bacterial adaptation driven by novel alleles on multicopy plasmids. To be specific, we will often refer to the evolution of antibiotic resistance throughout the manuscript. Yet, the results equally apply to other beneficial alleles. Using the mathematical theory of multitype branching processes, we determine the establishment probability of a novel allele that initially arises on a single plasmid copy within a single cell. Using these results, we calculate the probability of evolutionary rescue – i.e. the probability that the bacterial population escapes extinction following environment change through adaptive evolution – if the novel allele appears through mutation in an existing plasmid-carried gene. This probability depends on the mutational input and on the establishment probability of mutations. The plasmid copy number often has antagonistic effects on these two factors, making it non-obvious whether a higher or lower copy number leads to a higher probability of rescue. We especially explore how the relationship between the plasmid composition within a cell and bacterial fitness influences the results. As an example, we apply the framework to a simple model for antibiotic resistance through enzymatic antibiotic degradation, in which we derive this relationship mechanistically. We finally extend the modeling framework to account for adaptation from the standing genetic variation.

## The Model

We consider a bacterial population of initially *N*_0_ cells. Each bacterial cell contains exactly *n* copies of a non-transmissible plasmid. We distinguish two variants of this plasmid: the wild-type plasmid and the mutant plasmid. Initially, all cells are homozygous for the wild-type plasmid and sensitive to an antibiotic in their environment. The population size therefore declines due to the drug pressure, and the population will go extinct unless resistance evolves (see Fig. 1B). A resistance allele can appear through a single mutation of a gene on the multicopy plasmid (mutant plasmid). As explained above, this is possible if the wild-type plasmid carries a resistance gene that confers resistance to some antibiotic but not to the one currently present (see San Millan *et al*., 2016). The level of resistance of a cell depends on its plasmid composition. Further below, we extend our model to include standing genetic variation.

**Figure 1:**
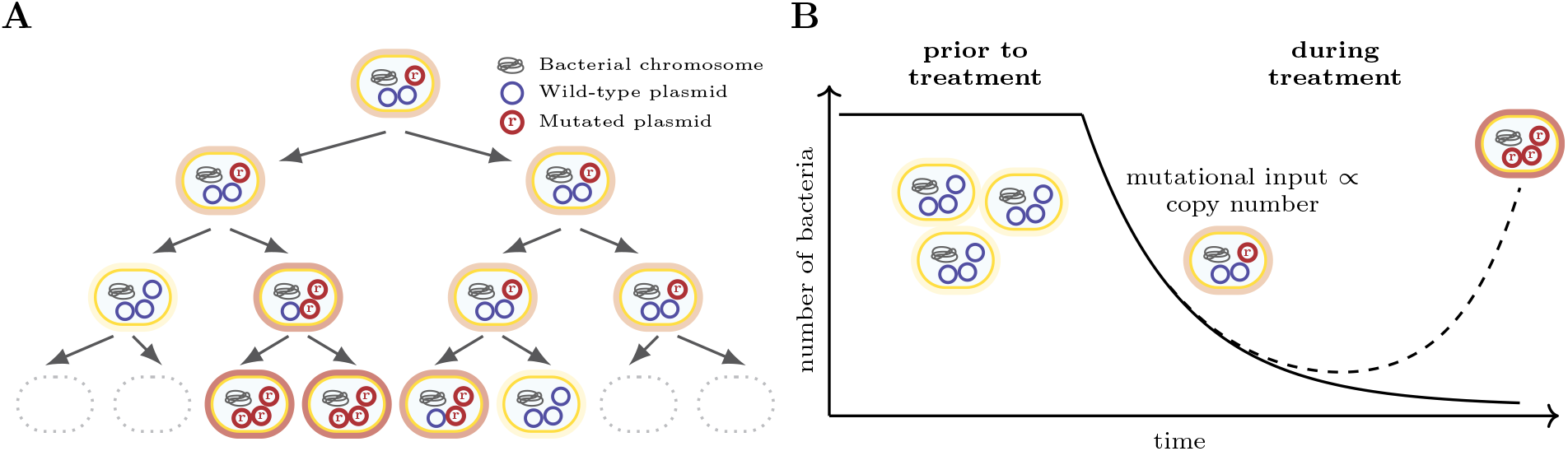
Bacterial adaptation on multicopy plasmids. Panel A: Segregation of plasmids at cell division and establishment of the resistance mutation (figure adapted from San Millan *et al*. (2016)). Panel B: Evolutionary rescue through *de novo* mutations on a multicopy plasmid.

We model the population dynamics by a birth-death process with birth (cell division) and death rates 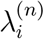 and 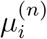 for a cell with *n* plasmids and *i* mutated plasmids. We denote the Malthusian fitness of a bacterial cell by 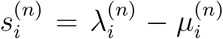. For cells that are homozygous for the wild-type plasmid (*i* = 0) and hence sensitive to the antibiotic, it holds that 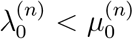. With an increasing number of mutant plasmids in a cell, the birth rate increases 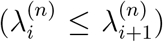, and/or the death rate decreases 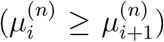. We normally assume that at least the cell type with only mutated plasmids (*i = n*) is resistant, so that 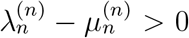. Otherwise the bacterial population could not escape ultimate extinction. The shape of 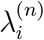 and 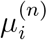 as a function of *i* reflects the dominance relationship between the mutant and the wild-type allele. Gene dosage effects can lead to a higher fitness of mutant homozygotes with increasing plasmid copy number (i.e. 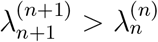 or 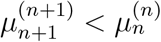). On the other hand, plasmids can impose a cost on the cell, and it is plausible to assume that this cost increases with the copy number. We implement plasmid costs as an increase in the death rate that depends linearly on *n*, e.g. 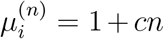. However, for most part, we ignore plasmid costs (*c* = 0) in order to disentangle the cost-independent effects of *n* on rescue. For all numerical examples presented in the figures, we choose 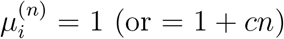 and 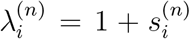. We set a hard carrying capacity of *N*_0_ cells. When the population size exceeds *N*_0_, a random cell is removed.

We assume that plasmid replication and cell division are perfectly synchronized. Right before cell division, each plasmid is replicated exactly once. At plasmid replication, mutation from the wild-type to the mutant plasmid occurs with probability *u* per plasmid. We neglect back mutations. Then the cell divides, and each daughter cell receives exactly *n* plasmids. The distribution of wild-type and mutant plasmids to the daughter cells, however, is random. The probability that a cell that contains *i* mutant plasmids before and 2*i* + *x* mutant plasmids after plasmid replication (*x* denotes here the number of newly mutated wild-type plasmids) divides into daughter cells with *k* and (2*i* + *x*) − *k* mutant plasmids, respectively, is given by

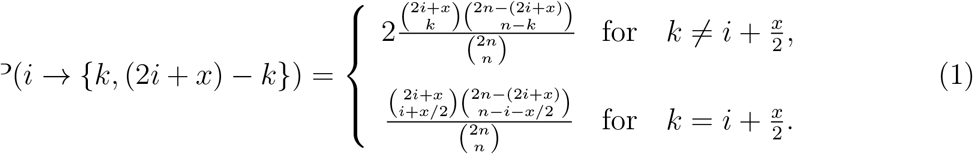

E.g., disregarding mutation (*x* = 0), a cell with two plasmids (*n* = 2), one of which is mutated (*i* = 1), divides into two heterozygous daughter cells with probability 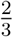 and in a homozygous wild-type and a homozygous mutant cell with probability 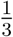.

The chosen model for plasmid replication and segregation is mathematically convenient and captures the essence of the process by introducing stochasticity in the number of mutant plasmids inherited by each daughter cell. However, it contains several simplifying assumptions. For example, plasmids are normally replicated after cell division. Thereby, some plasmids may be copied several times, and others not at all. An equal distribution of the plasmid copies contained in the mother cell to the daughter cells is an idealization, reflecting a perfect partitioning system. How segregation occurs varies across plasmids and moreover differs between low-copy and high-copy plasmids. Given the variation in segregation mechanisms and differences between plasmids with low and high copy numbers, no model fits them all, and we here choose one extreme. To test the robustness of our results, we set up and analyze two alternative models for plasmid replication and segregation in the supplementary information, section S1, in which we relax some of the assumptions.

## Analysis and Results

### Stochastic computer simulations

We perform stochastic computer simulations that exactly implement the model. The code is written in the C++ programming language, following the Gillespie algorithm (Gillespie, 1976) and making use of the GNU Scientific library (Galassi *et al*., 2009). We track the bacterial population until it has either gone extinct or until the mutant homozygote has reached a critical number *N_c_* above which the probability of extinction of the homozygote population is less than 1%. We determine this threshold mathematically through 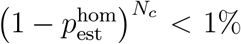, where 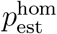 is the probability that a single mutant homozygous individual establishes a permanent lineage of offspring. We provide an expression for 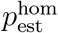 in Eq. (A.6).

### Mathematical approach

Whether the bacterial population successfully adapts or goes extinct depends on two factors: the mutational supply and the establishment probability of the resistance mutation once it has appeared. New resistance mutations appear approximately at rate 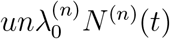 during population decline, where *N*^(*n*)^(*t*) is the number of wild-type homozygous cells at time *t*. (The approximation assumes that within any one cell, at most one plasmid copy acquires a mutation during replication.) The mutational supply hence linearly increases with the plasmid copy number *n*. Since the initial cell population is large, we describe the dynamics of wild-type homozygous cells deterministically (cf. Orr and Unckless, 2008; Tazzyman and Bonhoeffer, 2014; Uecker *et al*., 2014; Uecker and Hermisson, 2016; Uecker, 2017). Hence:

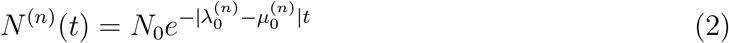

Yet, for evolutionary rescue to occur, it is not sufficient that a resistance mutation appears before the wild-type population goes extinct. It also needs to escape stochastic loss. We hence do not need to compute the rate of appearance of resistance mutations but the rate of appearance of *successful* resistance mutations. Any resistance mutation first appears in a single plasmid copy within a single cell, and we denote by 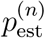 its establishment probability. Then, the rate of appearance of successful resistance mutations is given by 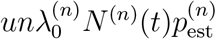. To determine 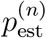, we use the mathematical theory of multitype branching processes (see Appendix A).

With this, the probability of evolutionary rescue by *de novo* mutations can be approximated by

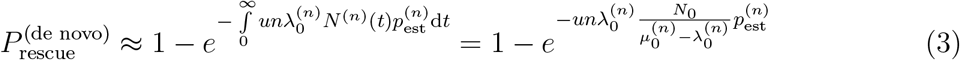

(see Orr and Unckless, 2008; Martin *et al*., 2013; Tazzyman and Bonhoeffer, 2014; Uecker *et al*., 2014; Uecker and Hermisson, 2016; Uecker, 2017; Anciaux *et al*., 2018). The exponential function is the zeroth term of a Poisson distribution and gives the probability that no successful resistance mutation appears in time, i.e. that the population goes extinct. Note that within this approach, rescue on a single copy plasmid is identical to rescue on a haploid chromosome (assuming equal mutation probabilities).

### Analytical solutions for one and two plasmid copies per cell

In a first step, we consider plasmids with copy numbers *n* = 1 and *n* = 2. For single-copy plasmids, establishment of the resistance mutation is simply determined by the dynamics of cell division and death, while for two-copy plasmids, segregation plays a role since heterozygous cells can either have two heterozygous or two homozygous daughter cells. The potential production of a daughter cell that is homozygous for the maladaptive wild-type allele *a priori* opposes establishment of the resistance mutation on a two-copy plasmid. On the other hand, gene dosage effects may outweigh this disadvantage. Moreover, the mutational input 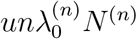 is twice as high on a two-copy plasmid (assuming that the plasmid copy number does not affect the fitness of wild-type homozygotes).

For *n* =1 and *n* =2, simple analytical solutions for 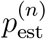 exist (from Eq. (A.2)):

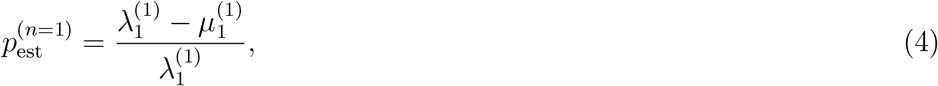

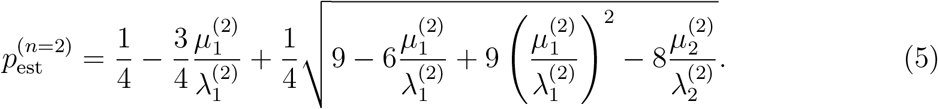

This allows us to analyze in detail how these effects play out. Under which conditions is the establishment probability of a beneficial allele higher on a two-copy than on a single-copy plasmid? And when is evolutionary rescue more likely?

We distinguish two cases: (1) There are no gene dosage effects, and the level of resistance depends on the *relative* number of mutated plasmids in the cell, hence 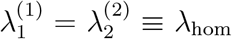 and 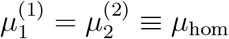. (2) There are gene dosage effects, and the level of resistance depends on the *absolute* number of mutated plasmids in the cell, hence 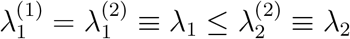 and 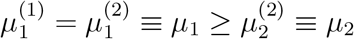. In both cases, we assume that there is no cost associated with a higher plasmid copy number.

It is intuitive and can also be derived from a comparison of the formulas for 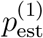 and 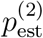 that in scenario (1), the establishment probability is always higher if the cell carries one plasmid than if it carries two. However, with two plasmid copies, the mutational input is higher, which may outweigh the lower establishment probability. Inserting formulas (4) and (5) into Eq. (3) and comparing the rescue probabilities shows that rescue is more likely to occur on a two-copy plasmid if

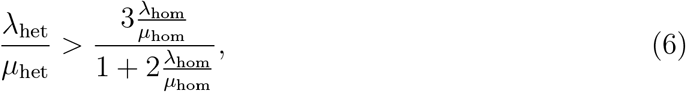

where λ_het_ and *μ*_het_ are the birth and death rates of the heterozygous two-copy cell (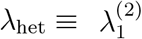 and 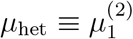). The fitness of heterozygous cells hence needs to be sufficiently high; otherwise the higher mutational input for *n* = 2 is insufficient to outweigh the lower establishment probability. In particular, evolutionary rescue is less likely on a two-copy plasmid than on a single-copy plasmid if heterozygous two-copy cells have a negative Malthusian fitness λ_het_ − μ_het_. In the special case 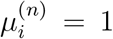 for all *n* and *i* and λ_hom_ = 1 + *s* and λ_het_ = 1 + *αs*, condition (6) simplifies to

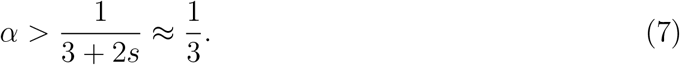

The variable *α* should not be confounded with the dominance coefficient of the mutation since the effect of the mutation is not *s* but 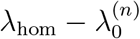. The dominance coefficient above which rescue is more likely on a two-copy than on a single-copy plasmid depends both on 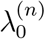 and on the beneficial effect of the mutation. The higher 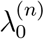 and the more beneficial the mutation, the lower is the required dominance coefficient.

In scenario (2) with gene dosage effects, the establishment probability can be higher with two plasmid copies than with one. This is the case if

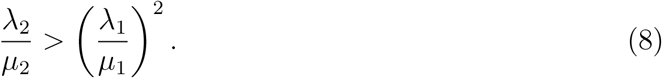

For the probability of evolutionary rescue, we find that under the above assumptions, two plasmid copies are always advantageous (see section S2 in the supplementary information). Remember though that we assumed that the second plasmid copy did not impose any additional cost. An additional cost, if strong enough, can make the second copy disadvantageous despite gene dosage effects.

### Arbitrary copy numbers

In the previous section, we saw that – as is also intuitively expected – the fitness of heterozygous cells and potential gene dosage effects determine whether a copy number of one or two is advantageous. Here, we proceed to investigate for arbitrary copy numbers how the relationship between the plasmid composition of a cell – i.e. the number of mutant plasmids *i* and wild-type plasmids (*n − i*) – and the cell’s fitness influences the form of *P*_rescue_(*n*). In the absence of gene dosage effects (i.e. if the fitness of mutant homozygotes is independent of *n*), we refer to this relationship as the dominance function of a mutant allele. In order to determine the establishment probabilities 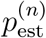, we solve Eq. (A.5) numerically (and then use Eq. (A.1)). Throughout this section, we set 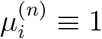 for all *n* and *i*, and only the birth rate 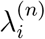 depends on *n* and *i*. Wild-type cells have a Malthusian fitness 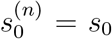 that is independent of *n*. We consider examples of three different types of plasmid mutations with different shapes of dominance functions – (i) dominant mutations (Fig. 2A), (ii) recessive mutations (Fig. 2B), (iii) mutations of intermediate dominance (Fig. 2C) – and (iv) mutations with a gene dosage effect (Fig. 2D). For (i)-(iii), 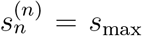 is independent of *n*, while it increases with *n* in case (iv).

For *n* = 1, the fate of the mutant plasmid is entirely determined by the birth and death dynamics of the mutant cell. In contrast, for higher copy plasmid numbers, segregation of mutant plasmid during cell division of heterozygous cells leads in general to one daughter cell with a lower number of mutant plasmids, stretching the establishment process of the mutant plasmid over a larger number of cell generations. If the fitness of mutant homozygotes is independent of *n* (cases (i)-(iii)), establishment of the mutant plasmid gets therefore less likely with increasing copy number *n* irrespective of the concrete dominance function (Fig. 2E-G). How strongly the establishment probability drops with *n*, however, depends on the dominance function.

For dominant mutations (case i), cells only need to carry one mutant plasmid to be resistant. Formally, dominant mutations are defined by 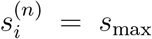 for *i* > 0. Then, 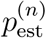 only moderately decreases with *n* and eventually converges to a constant (Fig. 2E). We find that the increase in the mutational supply with higher copy numbers exceeds the decrease in the establishment probability such that the probability of evolutionary rescue increases with the plasmid copy number (Fig. 2I).

For recessive plasmid mutations (case ii), only cells where all plasmid copies are mutated are resistant. Cells with a heterozygous plasmid composition are, just as wild-type cells, fully susceptible (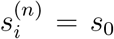 for *i < n*). Establishment probabilities for this plasmid type fall sharply with increasing plasmid copy numbers to very low values (Fig. 2F). Since heterozygous cells are susceptible to the antibiotic and are less likely to divide than to die, it is unlikely that a resistant homozygous mutant cell establishes if many cell divisions are required to generate it. Unlike for dominant mutations, rescue probabilities decrease with the plasmid copy number (Fig. 2J). The drop in the establishment probability is so dramatic that it cannot be compensated for by the higher supply of new mutations for higher plasmid copy numbers.

For intermediate dominance (case iii), we assume a Malthusian fitness 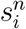 that increases linearly with the relative number of mutant plasmids, i.e. 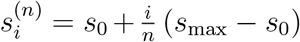. The behavior of 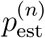 is in-between the two extremes of dominant and recessive mutations (Fig. 2G). If the fitness of homozygous mutant cells *s*_max_ is large, we observe a slight maximum in the rescue probability as a function of the plasmid copy number (Fig. 2K). This is the result of the antagonistic effects of the plasmid copy number on the establishment probability (decrease with *n*) and on the mutational input (increase with *n*). This maximum is shifted to higher plasmid copy numbers with increasing *s*_max_. If the fitness benefit of the mutation is small, *P*_rescue_ monotonically decreases with *n*.

Finally, we discuss the dependence of establishment and rescue probabilities on the plasmid copy number for mutations with a gene dosage effect (case iv). In this case, fitness of a bacterium depends on the absolute number of mutated plasmids within a cell, and cells with different plasmid copy numbers need the same absolute number of mutated plasmid copies to get resistant. We here implement a linear increase of the fitness with *n*, i.e. 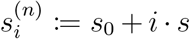. In that case, we find that the establishment probability increases with the plasmid copy number since the negative effect of segregational drift on establishment of the mutant plasmid is outweighed by the higher fitness of mutant homozygotes (Fig. 2H). In the limit of high copy numbers, the establishment probability reaches a constant value. If the increase in fitness per beneficial plasmid s is small and the plasmid copy number low, mutant homozygotes do not have a positive Malthusian fitness and the establishment probability of the mutant plasmid is hence zero (see inset of Panel H). Obviously, since both the mutational input and 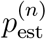 increase with *n* under the implemented gene dosage effects, rescue is more likely for high than for low copy number plasmids (Fig. 2L).

In the supplementary information, section S1, we show the rescue probabilities under two alternative models of plasmid replication and segregation and compare the results with those presented here. General trends are robust. Yet, especially for recessive mutations, differences appear. The decline of *P*_rescue_ with *n* is much less strong under the alternative schemes, and low but intermediate copy numbers may even be best (though only by a very slight margin) for the bacterial population.

**Figure 2:**
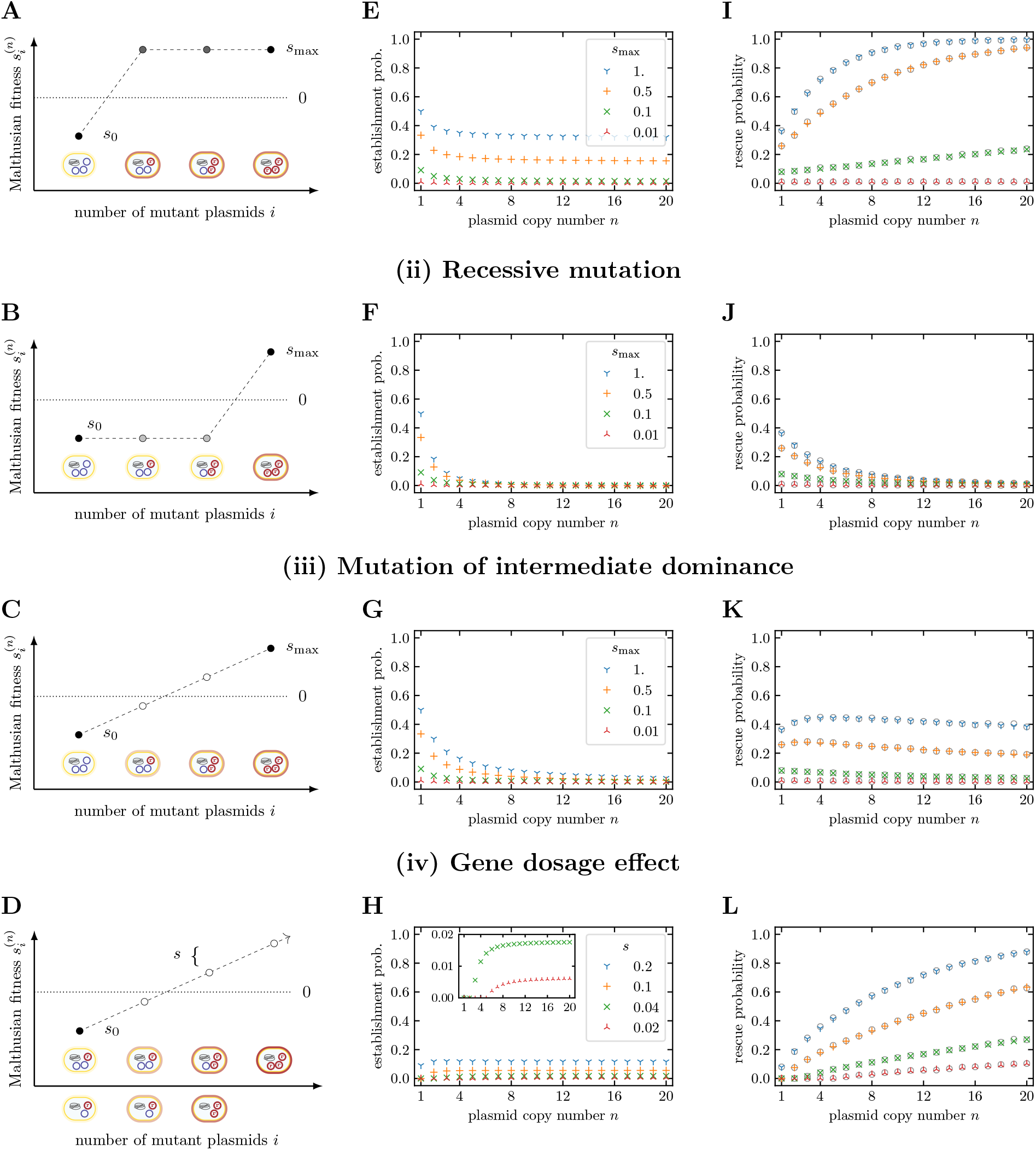
The influence of the dominance function of a plasmid mutation and of gene dosage effects on the establishment of resistance-conferring plasmid variants and the probability of evolutionary rescue. We consider three generic types of dominance functions for plasmid mutations (i-iii) and gene dosage effects (iv). Panels A-D: Illustration of the relation between cell types and their Malthusian fitness. Panels E-H: Establishment probabilities of a plasmid mutation arising on a single plasmid copy. Panels I-L: Probability of evolutionary rescue from *de novo* mutations. Without gene dosage effects, the establishment probability decreases with the plasmid copy number. Yet, the probability of evolutionary rescue may still increase with *n* if the decline in the establishment probability is compensated by the increased mutational supply. Results for the establishment probabilities were obtained numerically by using Eq. (A.5) with Eq. (A.1). These were inserted into Eq. (3) to obtain the probability of evolutionary rescue. Parameters: *s*_0_ = −0:1, *uN*_0_ = 0.1, *N*_0_ = 10^5^. Open circles show averages over 104 stochastic simulations.

### Example: The influence of the antibiotic concentration

So far, we have fixed the relationship between the plasmid composition of a cell and its fitness. Yet, this relationship derives from the specific resistance mechanism and likely from the environmental conditions. Here we use a simple model to demonstrate how the above formalism can be applied to concrete situations of resistance evolution. Using a basic mechanistic model for cell resistance, we establish how the dominance function depends on the external antibiotic concentration to which the bacterial population gets exposed. We then determine the concentration-dependent probability of evolutionary rescue for a range of plasmid copy numbers.

We assume that the fitness of a bacterial cell depends on the antibiotic concentration *c*_in_ within the cell and take the corresponding dose-response curve *S*(*c*_in_) as given (Fig. 3A and Eq. (B.1)). For the wild-type, the internal concentration *c*_in_ equals the antibiotic concentration in the environment *c*_out_. A decrease of the internal concentration occurs through enzymes that degrade the antibiotic within the cell (Eq. (B.3)). The corresponding gene is located on the plasmid and carries a mutation in the mutant plasmid. The mutation does not impose a fitness cost. While both plasmid variants, wild-type and mutant plasmids, code for the enzyme, only the mutated version can degrade the specific antibiotic (Fig. 3B). The rate of antibiotic degradation increases with the number of mutated plasmids within the bacterial cell. Consequently, the internal concentration decreases with *i*, and bacterial fitness increases as determined by the shape of the dose response curve (Fig. 3A). Details on the model can be found in Appendix B.

We distinguish two limiting cases. In the first case, we assume that the resources (e.g. proteins) needed for the production of the degrading enzyme are limited. The limiting factor that determines the rate of enzyme production is not the number of mutated plasmid copies but the available resources for which wild-type and mutant plasmids are competing. In that case, the rate of mutated enzyme production and thus the antibiotic degradation rate depend on the relative number of mutated plasmids *i/n* per cell (see the inset of Fig. 3C for the fitness of different cell types as a function of *c*_out_). Concretely, we assume that the enzyme production increases linearly with *i/n* (Eq. (B.4)). We find that the optimal plasmid copy number *n* changes with the antibiotic concentration (Fig. 3C). The reason is that the dominance function changes along the antibiotic gradient (Fig. 2D). For low concentrations, a small change in the internal antibiotic concentration (*i/n* small) leads to a large fitness increase (Fig. 3A) such that the mutation has a concave dominance function (*c*_out_ = 1.1 in Fig. 3D). This resembles the dominant mutation from the previous section. For these concentrations, a high plasmid copy number maximizes population persistence. For high concentrations, in contrast, a high fraction of mutant plasmids is required to achieve an appreciable increase in fitness (Fig. 3A). Then, the dominance function is convex, resembling a recessive mutation (*c*_out_ = 1.6 in Fig. 3D). In this regime, a low copy number leads to the highest probability of evolutionary rescue. For intermediate concentrations, the dominance function has an inflection point and an intermediate copy number is most advantageous for the bacterial population (*c*_out_ = 1.3 and *c*_out_ = 1.4 in Fig. 3).

In the second case that we consider, the enzyme production and the degradation rate depend on the absolute number of mutated plasmids *i*, i.e. not the resources needed for enzyme production are the limiting factor but the number of mutant plasmids. This reflects a gene dosage effect. We assume here that the enzyme production increases linearly with *i* (Eq. (B.5)). The Malthusian fitness of cell types with *i* = 0, 1, 2, … mutated plasmid copies as a function of the external antibiotic concentration is shown in the inset of Fig. 3E, assuming as in the previous section that plasmids do not impose a fitness cost. At all concentrations, rescue is more likely for higher *n* (Fig. 3E).

This simple picture changes if plasmids have a high fitness cost, which increases the death rate by *c* per copy number in our model. In that case, the fitness of wild-type cells decreases with the plasmid copy number, and even for mutant homozygotes, a higher copy number does not always imply a higher fitness (see the inset of Fig. 3F, where dashed lines indicate the wild-type and straight lines the mutant). In contrast to the results without plasmid costs (Panel E) where the highest copy number always led to the highest chance of evolutionary rescue, the optimal copy number now depends on the antibiotic concentration. For low concentrations, the high burden imposed by a high copy number leads to a low probability of rescue. However, for higher concentrations, where efficient antibiotic degradation is important, rescue is more likely with higher plasmid copy numbers even though carrying plasmids is costly.

### The mutant frequency in the standing genetic variation

So far, we have focused on evolutionary rescue from *de novo* mutations that arise during the decline of the bacterial population after the change in the environment, i.e. in the presence of antibiotics. Yet, mutated plasmids may be present in the population prior to environmental change. Here, we extend our model to account for rescue from the standing genetic variation. We assume that mutant plasmids that confer resistance in the antibiotic environment are disadvantageous in the absence of antibiotics and segregate in mutation-selection balance in the population. We denote by −*σ* the selection coefficient for mutant homozygous cells for mutations, where the relative proportion of mutated plasmids determines the cell fitness, and by −*σ_GDE_* the deleterious effect per plasmid copy if there is a gene dosage effect. To determine the frequency of mutant plasmids in the absence of antibiotics, we adapt a multitype Moran model that describes the dynamics of the various cell types in a bacterial population of constant size 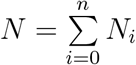, where *N_i_* denotes the number of cells with *i* mutated plasmid copies. From the Moran model, we derive a set of ordinary differential equations, from which we numerically obtain the numbers of mutant homozygous and heterozygous cells in deterministic mutation-selection equilibrium. From those, we estimate the contribution of the standing genetic variation to the probability of evolutionary rescue. Since the population faces ultimate extinction if all resistance mutations go extinct, the rescue probability can be approximated by

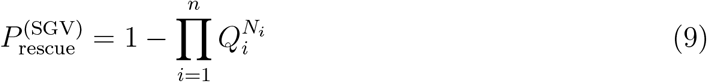

where *Q_i_* denotes the probability that a cell of type *i* does not leave a permanent lineage of descendants. A detailed description of the model is given in Appendix C. The deterministic approach assumes an infinite population size and neglects stochastic fluctuations in *N_i_*. We therefore additionally perform simulations of the stochastic model to account for finite populations.

**Figure 3:**
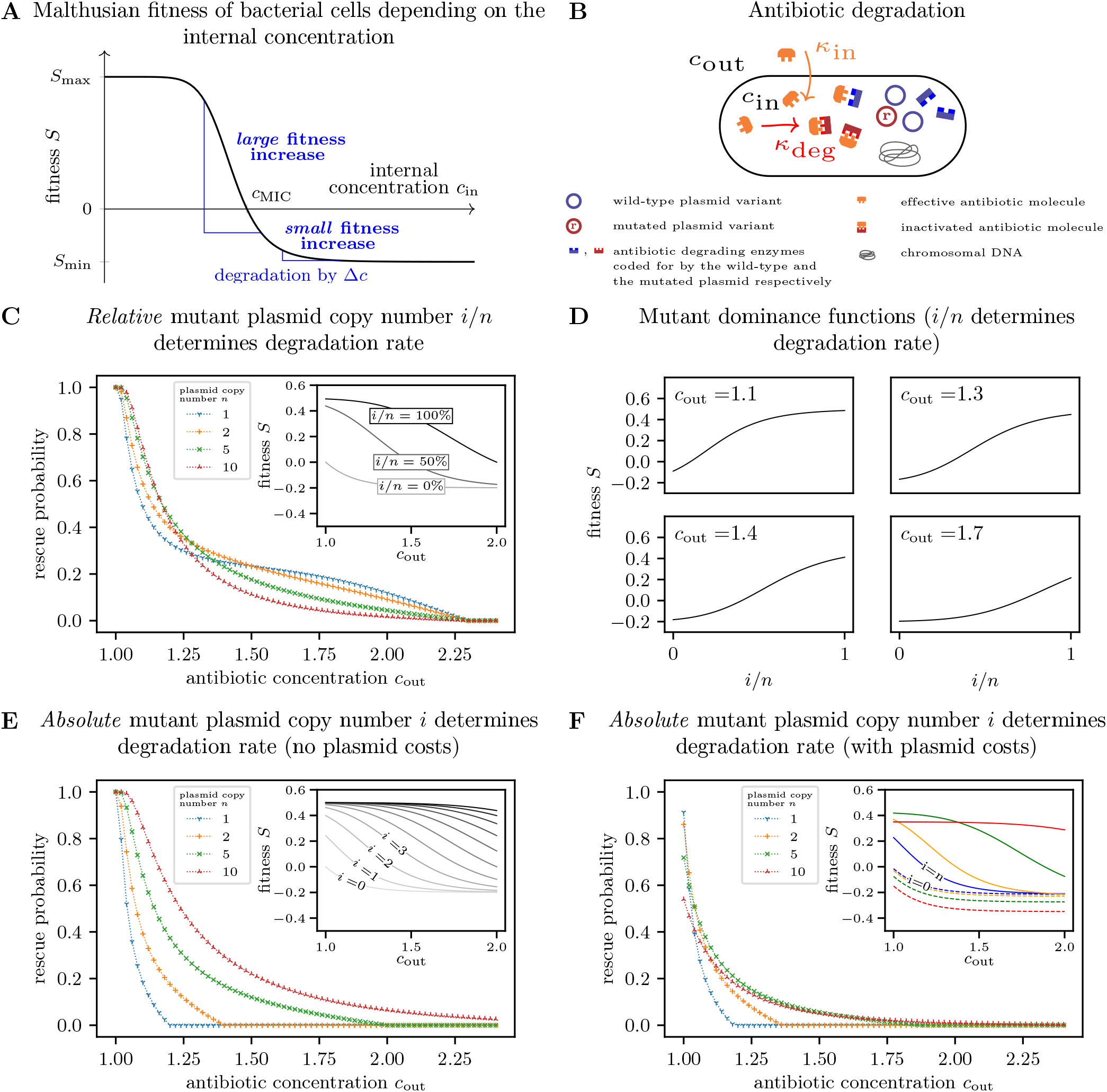
Dependence of the optimal plasmid copy number on the antibiotic concentration. Panel A: Malthusian fitness *S* of a bacterial cell as a function of the antibiotic concentration within the cell. Panel B: Illustration of a simple model to determine the antibiotic concentration within the cell. Antibiotic molecules diffuse into the cell, where they get inactivated due to reactions with the mutant version of a plasmid-encoded enzyme. We assume that inflow of antibiotic molecules *κ*_in_ and degradation *κ*_deg_ are in equilibrium. Panel C: Probability of evolutionary rescue if the relative number of mutant plasmids within the cell determines the degradation rate. The enzyme production increases linearly with the relative number of mutated plasmid copies, and the cell fitness is thus determined by the fraction *i/n* of mutated plasmids within the cell (inset). Panel D: Mutant dominance functions for various antibiotic concentrations if the relative number of mutant plasmids i/n within a cell determines the degradation rate. With increasing concentration, the shape changes from concave to convex. Panel E: Probability of evolutionary rescue if the absolute number of mutant plasmids within the cell determines the degradation rate. The enzyme production increases linearly with the absolute number of mutated plasmid copies, corresponding to a gene dosage effect (see inset for the concentration-dependent fitness of each cell type). Rescue probabilities increase with higher plasmid copy numbers. Panel F: Probability of evolutionary rescue if the absolute number of mutant plasmids within the cell determines the degradation rate and plasmids impose a copy-number-dependent cost. Unlike in Panel E, plasmid costs increase bacterial death rates by *c* = 0.015 per plasmid copy. For very low antibiotic concentrations, the probability is lowest for high copy numbers due to the high associated costs. For high concentrations, the disadvantage of a higher burden is outweighed by the advantage of more efficient antibiotic degradation, and the rescue probability increases with the copy number. Results for the rescue probabilities were obtained as in Figure 2. The fitnesses are calculated from Eq. (B.1) and Eq. (B.7) with either Eq. (B.4) (for Panel C and D) or Eq. (B.5) (for Panel E and F). Parameters: *N*_0_*u* = 0.2, *S*_max_ = 0.5, *S*_min_ = −0.2, *c*_MIC_ = 1, *κ* = 8, *η* = 1.3, *ω* = 0.2, *γ* =1.

Before considering rescue probabilities, we briefly discuss the distribution of the frequencies of the various cell types in the standing genetic variation, comparing dominant and recessive mutations. Figure 4 shows, in an exemplary manner for a bacterial population with *n* =10 plasmid copies per cell, the expected frequencies *N_i_/N* for different strengths of selection against the mutant plasmid. As expected, the frequency of cells containing mutated plasmids is higher for a recessive than for a dominant mutation, since in that case the mutation is only exposed to selection if it occurs in a mutant homozygous cell (cf. also Fig. 5A and B). Since cells with a heterozygous plasmid composition are not subject to selection for recessive mutations, their frequencies do not visibly change with the strength of selection. The frequency of homozygous mutant cells, however, strongly increases with decreasing purifying selection, and mutant homozygotes become the most frequent mutant class for weak selection. For weak selection against the mutant plasmid, the difference between recessive and dominant mutations mainly shows in classes with high numbers *i* of mutant plasmids and most prominently in the number of mutant homozygotes. Generally, the variance in the cell frequencies is high (see error bars in Fig. 5 and Fig. S3.4). We will discuss consequences of this high variance below.

As for rescue from *de novo* mutations, we are interested in how the probability of rescue from the standing genetic variation 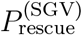 depends on the plasmid copy number *n*. For this, we assume that the dominance function remains the same upon the environmental change, e.g. mutations that are dominant prior to the change are also dominant afterwards. Likewise, gene dosage effects persist such that a mutant homozygous cell is particularly fit in the new environment but particularly unfit in the old environment.

Fig. 5 shows the expected cell type frequencies, the probability of rescue from the standing genetic variation, and the total rescue probability as a function of the plasmid copy number. Both for dominant and for recessive mutations as well as for mutations of intermediate dominance, the expected overall frequency of mutant cells 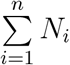 increases with the plasmid copy number *n* due to the increased mutational supply (Fig. 5A-C). The effect is particularly strong for recessive mutations, where the deleterious mutation is masked in heterozygous cells. For recessive mutations, we moreover observe that the frequency of mutant homozygotes *N_n_* is (nearly) independent of the plasmid copy number (Fig. 5B). Generally for high *n*, irrespective of the dominance function, most cells only harbor few (less than 50%) mutated plasmids. Especially for recessive mutations or those with intermediate dominance, these cells are not very likely to leave a permanent lineage of descendants after the environmental change. Therefore, the probability of evolutionary rescue increases only slightly (if at all) with *n* (Fig. 5E-G). If there is a gene dosage effect, i.e. if selection against the mutation increases with the absolute number of mutant plasmids, there is a minimum in the frequency of mutant cells for intermediate plasmid copy numbers *n*, stemming from the antagonistic effects of the plasmid copy number on the mutational supply and on the strength of selection against the mutant plasmid (Fig. 5D). Yet, since the establishment probability strongly increases with *n*, rescue from the standing genetic variation still increases with the plasmid copy number for the chosen parameter set (Fig. 5H). Further analysis shows that, if the mutation is highly beneficial after the environmental change, a slight minimum in 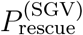 as a function of *n* may exist (e.g. for λ_0_ = 0.95, *σ_GDE_* = 0.1, *s* = 2, *N* = 3 × 10^8^, *u* = 3 × 10^−10^).

Unless the selective advantage of the mutant plasmid is very weak after the environmental change, the deterministic approximation for *N_i_* leads to an overestimation of the rescue probabilities from the standing genetic variation (see Fig. 5E for *s*_max_ ∈ {0.1,0.5,1} and Panel H for *s* ∈ {0.04, 0.1, 0.2}). The error gets larger for smaller population sizes (Fig. S3.3). The reason for the deviation between the analytical theory and the stochastic simulations is the large variance in *N_i_*. Since 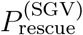 is strongly concave in *N_i_* for strong selection, the negative and positive effects of fluctuations in *N_i_* below or above the average do not cancel. Due to these stochastic effects, in Fig. 5F (recessive mutation), the probability of evolutionary rescue decreases for high copy numbers in contrast to the deterministic approximation, which predicts a monotonic increase; the same can be observed for mutations of intermediate dominance (Fig 5G with *s*_max_ = 0.5). It seems plausible that for high *n*, stochastic fluctuations in the numbers of cells with high *i* are stronger, causing this decrease. To probe this intuition, in Appendix S3, we investigate this behavior in some more detail for recessive mutations and a lethal wild-type in the new environment. In that case, rescue entirely relies on mutant homozygotes in the standing genetic variation. *P*_rescue_ then decreases with *n* if stochasticity in *N_n_* is taken into account but remains constant under the deterministic approximation of *N_n_* (Fig. S3.5). Nevertheless, in most cases, the trend of 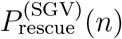 is correctly estimated based on the deterministic approximation of the cell type frequencies in the standing genetic variation.

Finally, we combined rescue from the standing genetic variation and from *de novo* mutations to determine the total probability of evolutionary rescue (Fig. 5I-L):

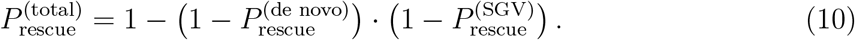

For mutations, where the relative number of mutant plasmids determines the cell fitness (cases (i)-(iii)), the effect of the dominance function on total rescue probabilities is much weaker than its effect on rescue by *de novo* mutations alone. In all three cases, rather similar total rescue probabilities are observed.

**Figure 4:**
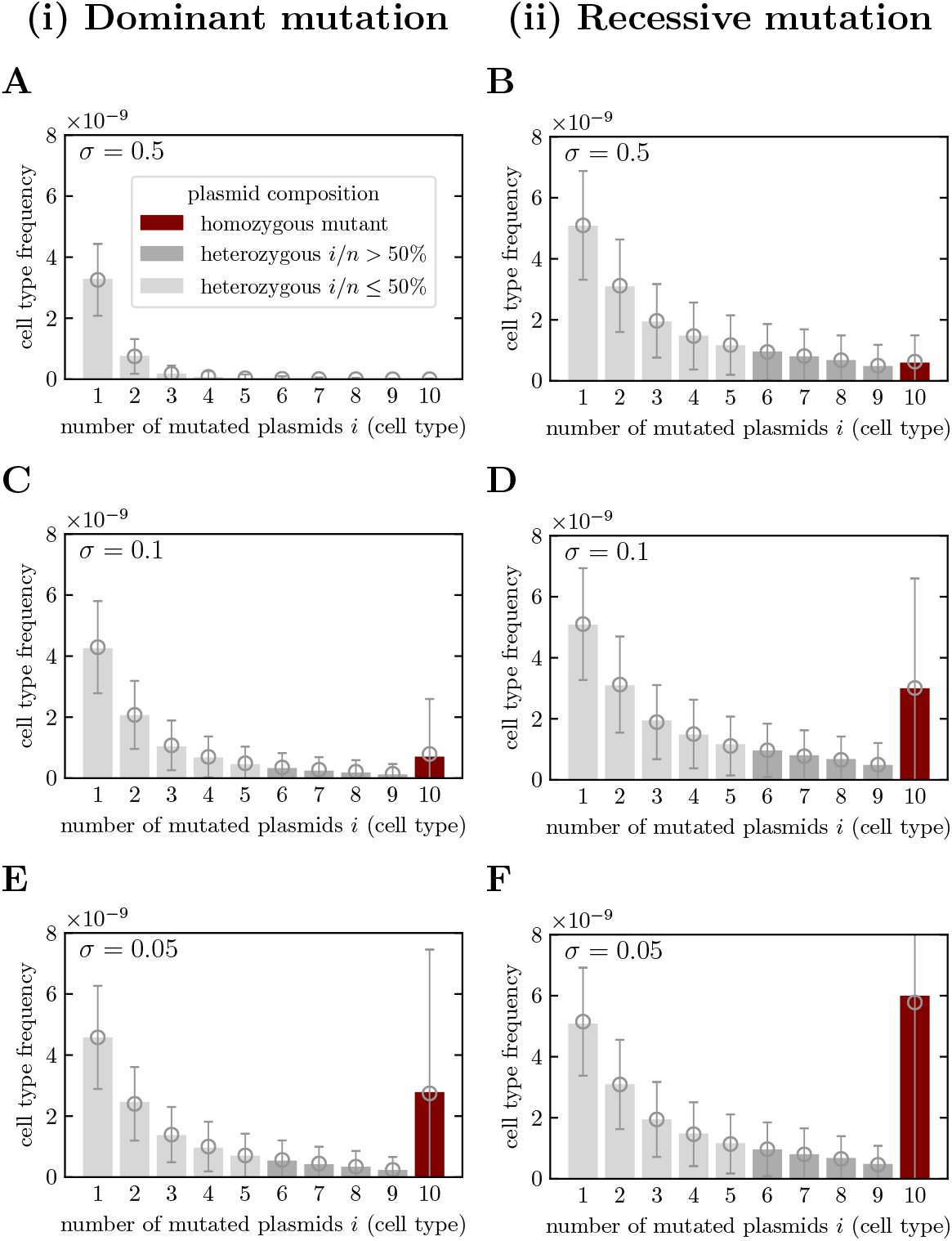
Distribution of mutant cell type frequencies in the standing genetic variation for dominant and recessive mutations. Mutant plasmids are deleterious prior to antibiotic treatment with a selection coefficient of −*σ* (see appendix C). Frequencies are obtained from the equilibrium of the deterministic model, i.e. by numerically integrating ((C.5), (C.6) and (C.7)). Parameters: *n* = 10, *N*_0_ = 3 × 10^9^, *u* = 3 × 10^−10^. Open markers and error bars show averages and standard deviations of 10^3^ stochastic simulations.

## Discussion

Adaptation fundamentally depends on two factors, the presence of adaptive alleles and their probability to escape stochastic loss while rare. For plasmid-carried genes, the plasmid copy number influences both of these factors. On one hand, the mutational target size increases with the number of plasmid copies per cell. On the other hand, unlike for alleles on haploid genomes, it takes several cell divisions, until a cell with a large fraction of mutant plasmids is generated, negatively affecting the mutant plasmid’s chances to establish. We here developed a framework to model bacterial adaptation driven by novel alleles on multicopy plasmids. We determined the probability of evolutionary rescue, taking into account new mutations and mutations from the standing genetic variation.

**Figure 5:**
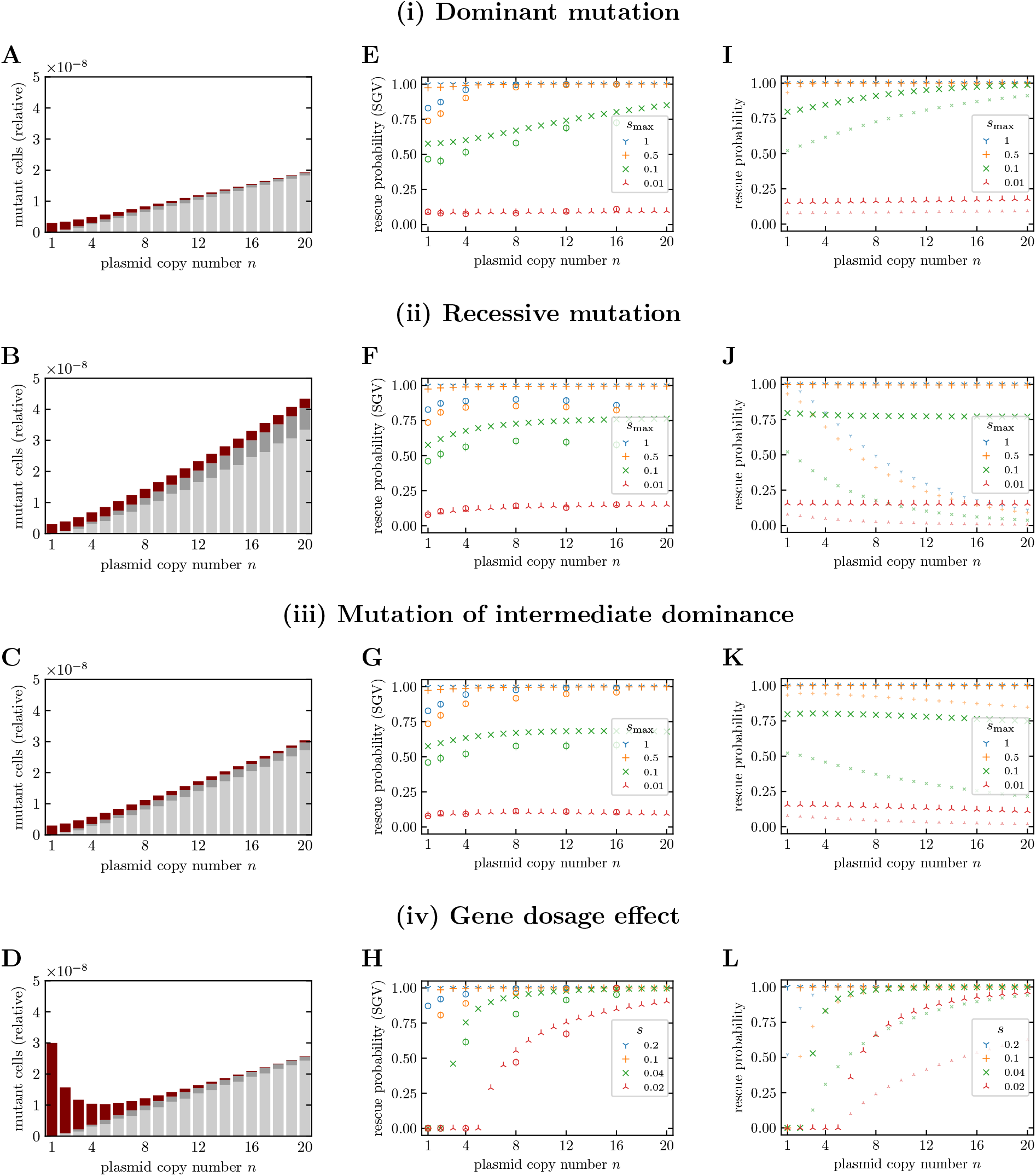
Dependence of mutant frequencies on the dominance function and the influence of standing genetic variation on the probability of evolutionary rescue. Panels A-D: Frequencies of mutant cells for a bacterial population in the standing genetic variation (SGV) for a range of plasmid copy numbers. Mutant plasmids are deleterious with a selection coefficient of −*σ* = 0.1 (dominant, recessive, intermediate). For the gene dosage effect (iv), each mutant plasmid decreases the selective fitness of its host cell by *σ*_GDE_ = 0.01. Panels E-H: Probabilities of evolutionary rescue from standing genetic variation (SGV) for those bacterial populations in an antibiotic environment. Panels I-L: Total probabilities of evolutionary rescue, including rescue from standing genetic variation and rescue from *de-novo* mutations. For comparison, small markers indicate rescue probabilities from *de-novo* mutations only. After the switch to the antibiotic environment, wild-type cells have a birth rate 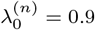. The fitness of resistant bacteria is increased by Smax for (i-iii) and by *s* per mutated plasmid for (iv) (analogous to Fig. 2). Frequencies (Panel A-D) are obtained from the equilibrium of the deterministic model, i.e. by numerically integrating Eqs. (C.5), (C.6) and (C.7). Rescue probabilities from SGV (Panels E-H) are obtained from Eq. (9) and Eq. (A.5). Total rescue probabilities including *de-novo* mutations (Panels E-H) are obtained from Eq. (10). Other parameters: *u* = 3 × 10^−10^, *N* = 3 × 10^9^. Open markers show averages over 10^3^ stochastic simulations. Error bars indicate twofold standard errors.

### The influence of the dominance function

It is no surprise that the dominance of the mutant allele is key for whether a low or a high copy number maximizes the probability of evolutionary rescue. We studied examples of dominant and recessive mutations and mutations of intermediate dominance and mutations in genes with a gene dosage effect, always assuming that bacterial fitness is increasing (or at least non-decreasing) with the number of mutated plasmid copies. Only for the latter, both the mutational input and the establishment probability increase with the plasmid copy number, while in all other cases, the effects are antagonistic. I.e., unless there are gene dosage effects, the plasmid copy number augments the mutational input but decreases the establishment probability. We observe that for dominant mutations, the benefits of a high per cell mutation rate outweigh the negative effects on the establishment probability such that the rescue probability increases with the plasmid copy number. This holds true both for *de novo* rescue and for rescue from the standing genetic variation. For recessive mutations, there is a phenotypic delay of several generations until the allele can be picked up by selection. The establishment probability therefore drops much more strongly with the plasmid copy number than for dominant mutations. In that case, the probability of *de novo* rescue decreases with *n*. On the other hand, in an environment where the allele is deleterious, its effect is masked in all heterozygous cells, leading to a high number of mutant cells in the standing genetic variation. In a deterministic analysis of the standing genetic variation, this leads to an increase in the probability of rescue from the standing genetic variation with *n*, even for recessive mutations (for consequences of stochasticity see below). In sum, for recessive mutations, the total probability of rescue remains constant as *n* increases.

These results are in line with the findings by Sun *et al*. (2018) for a model of bacterial adaptation on the chromosome. While bacteria are usually treated as haploid, making dominance inconsequential, Sun *et al*. (2018) take into account that bacterial chromosomes can be polyploid during growth such that the dominance of mutations strongly affects the adaptive process. The segregation mechanism is different from the one in the present model, since the chromosome copies are not randomly distributed to the daughter cells at cell division. Rather, all chromosomes carrying the mutation go into the same daughter cell. That both models make consistent predictions shows that the results are robust to the segregation mechanism (but see also the discussion further below). Indeed, we study two alternative models for plasmid replication and segregation, and these mostly confirm the conclusions except for a very slight maximum in the rescue probability for low copy numbers if the mutation is recessive.

Based on a deterministic analysis of the cell numbers in the standing genetic variation, both the present study and Sun *et al*. (2018) find that the mean number of mutant homozygous cells is independent of the plasmid copy number for recessive mutations. Stochastic simulations confirm that this holds for their mean number. Yet, the variance is large and increases with n. Therefore, the probability of rescue from the standing genetic variation starts to decrease with large copy numbers, an effect that was not previously observed.

In all our examples, we assume that the dominance function remains the same across environments. But for diploid organisms, for which dominance has been studied more intensely, it has been found that the dominance coefficient can be different in different environments (Gerstein and Otto, 2014). Indeed, within a simple model of antibiotic degradation, we find that the dominance function is concentration – and hence environment – dependent.

### Empirical dominance relationships

In this latter model, we derive the dominance relationship between mutated and wild-type plasmids mechanistically. In all other parts of the manuscript, however, we treat the dominance of mutations as a parameter. This raises the question about the nature of naturally occurring resistance determinants, and we here discuss a few examples that represent (partially) dominant or (partially) recessive mutations or resistance genes with gene dosage effects. Macrolide antibiotics such as erythromycin bind to the 23S RNA, ultimately interferring with protein synthesis (Vester and Douthwaite, 2001). Mutations in *rrn* genes, altering the ribosomal RNA, can confer resistance but these mutations have low dominance. Therefore, a sufficiently high fraction of gene copies need to be mutated to yield a resistant phenotype (Sigmund and Morgan, 1982; Mark *et al*., 1983; Weisblum, 1995; Sander *et al*., 1997; Vester and Douthwaite, 2001). Consequently, bacterial species that carry only one or two copies of the respective gene on the chromosome such as *Helicobacter pylori* or *Mycobacterium smegmatis* can acquire resistance through mutations in *rrn* genes while this is not possible in *Escherichia coli* that carries seven copies of the *rrn* operon (Sander *et al*., 1997; Vester and Douthwaite, 2001). Yet, Sigmund and Morgan (1982) isolated mutations conferring erythromycin resistance in *rrn* genes on a multicopy plasmid and demonstrated that *E. coli* carrying the mutant plasmid were resistant. Interestingly, Sigmund and Morgan (1982) discuss the possibility of resistance evolution via this pathway (i.e. *de novo* mutation of a *rrn* gene on a multicopy plasmid, followed by segregation to reach a high fraction of mutated copies in the cell). In the examples in our article, we do not consider genes that are present both in the chromosome and on the plasmid, e.g. essential genes such as the *rrn* genes that are additionally carried on a plasmid. This would lead to a more complex function for the type-dependent bacterial fitness that would depend on the number of chromosomal gene copies. While we did not explicitly study such a case, it is straightforward to implement. While mutations in *rrn* genes are partially recessive, mutations in loci that affect methylation of rRNA have been shown to be dominant (Heiser and Davies, 1972; Weisblum, 1995; Vester and Douthwaite, 2001). As another example, let us consider resistance to trimethoprim, an antibiotic that prevents DNA synthesis by binding to an enzyme (dihydrofolate reductase) in the DNA synthesis pathway. Resistance to trimethoprim is conferred by a plasmid carrying a mutated *drf* gene that codes for a mutated enzyme to which the antibiotic cannot bind and therefore provides a by-pass (Hall and Partridge, 2004; van Hoek *et al*., 2011). This indicates at least a certain degree of dominance. Last, as already pointed out, some resistance genes display a gene dosage effect. This is especially true for genes coding for antibiotic-degrading enzymes such as *β*-lactamases (Martinez and Baquero, 2000; San Millan *et al*., 2016). Noteworthily, there also exist negative gene dosage effects: it has been found that *E. coli* displays lower levels of resistance to tetracycline if the resistance-conferring transposable element Tn10 is carried on a multicopy plasmid than if it is present in a single copy on the chromosome (Moyed *et al*., 1983). Note also that in all the examples that we discussed here (except maybe for the last one), new resistance could occur through mutation in existing genes that can be plasmid-encoded and are hence relevant to the present study.

### Plasmid replication and segregation

We developed a mathematical framework to describe the dynamics of multicopy plasmids that allows for a clear and easy separation of the mutational input and the establishment probability of mutant alleles. We apply this model across the entire range of copy numbers. Obviously, this neglects large parts of the complexity of plasmid dynamics. We here briefly discuss some of the assumptions.

In the model presented in the main text, plasmid replication is initiated immediately before the host cell divides and each plasmid copy replicates once. In reality, plasmids replicate throughout the cell cycle (Bogan *et al*., 2001; Nordström, 2006). Fully accounting for this would require to use a full nested model that describes both the within-cell dynamics of plasmids and the birth-death dynamics of bacterial cells. This in turn would require knowledge of the period during the cell cycle during which the drug is effective, the number of plasmids that are present during that period, and the level of resistance they confer. Such a nested model would be appropriate to describe a specific system in detail but is less suitable to serve as a general model.

A more crucial assumption of our model of plasmid replication is that each plasmid copy reproduces exactly once. While (ideally) every plasmid replicates once per cell cycle *on average*, individual plasmid copies are picked for replication at random from the pool of all existing copies (Nordström, 2006). This implies especially that a cell with more than one mutated plasmid copy can appear from one cell generation to the next. We therefore also tested alternative models where plasmid replication is carried out iteratively (see supplementary information section S1). In these alternative models, we moreover inverted the order of plasmid replication and segregation. A comparison with the main text model showed that the trends in the probability of evolutionary rescue are mostly similar independent of the assumed scheme of plasmid replication.

The second set of assumptions concerns the segregation of the plasmid copies to the daughter cells. For the main text model, we assume that each daughter cell receives the exact same number of plasmid copies. Most low copy number plasmids possess active partitioning systems that force plasmid copies within a bacterial cell to opposite directions, leading to two (or few) clusters and at least partially balancing plasmid numbers in both daughter cells (Million-Weaver and Camps, 2014). High copy number plasmids do not possess any known active partitioning systems, although their existence has been discussed (Wang, 2017; Million-Weaver and Camps, 2014). Indeed, active partitioning systems are in that case less relevant, since with increasing copy number, it gets less and less likely that the plasmid gets lost even for a random distribution of plasmid copies to the daughter cells. Early models for the segregation of high copy number plasmids assume that the plasmids diffuse freely within the cell cytoplasm and are randomly distributed in space (Summers, 1991). However, it has later been found that high copy number plasmids form clusters within the cell (e.g. Pogliano *et al*., 2001). To explain the random distribution of high copy number plasmids to the daughter cells despite of clusters, it has been hypothesized that for replication, single plasmids get detached from the clusters and transported to the mid-cell, where they get replicated; afterwards, the two sibling plasmids diffuse in space and attach to existing clusters or become the founders of new ones (Nordström and Gerdes, 2003). Another (compatible) explanation is the observed existence of single plasmid copies in addition to the clusters (Wang, 2017). Assuming a random distribution of high copy number plasmids to the daughter cells, the relative deviation from an equal share decreases with the copy number. Overall, our assumption of an equal split of all plasmids seems to be a reasonable approximation for a baseline model. For one of the alternative models (no. 2) in the supplementary information, we have loosened the restriction of equally distributed plasmids.

The most critical assumption in all our model variants is the random distribution of wild-type and mutant plasmids at cell division. For high copy number plasmids, this assumption is generally considered as true (San Millan *et al*., 2016; Ilhan *et al*., 2019). Yet, for low copy number plasmids with active partitioning systems, it may not be well justified. The common view is that plasmids are recruited for replication to the center of the cell; afterwards, in the presence of active partitioning systems, the two replicates are pushed (or pulled) towards the two cell poles, thus separating them such that at cell division, they will end up one in each of the daughter cells (Nordström and Gerdes, 2003). Hence, in this case, our assumption might distort the chance of resistance evolution. If mutations are recessive, for example, a random distribution allows for an efficient accumulation of mutated plasmids, which is needed to generate phenotypically resistant cells. With a lack of random distribution for individual plasmid clones, homozygous mutant cells (resistant) are generated at a lower rate, leading to a lower chance of resistance evolution. The opposite effect might happen for dominant mutations, leading to a higher chance of resistance evolution due to the active partitioning of mutated plasmids to both daughter cells. While a deterministic partitioning of replicate plasmids is often proposed, there is also some evidence that partitioning could be more random. First, to explain the instability of different plasmid types with the same partitioning system, it has been suggested that heterologous plasmid pairs are formed in the cell center before partitioning, which would require a time delay between plasmid replication and partitioning (Nordström, 2006). For our model, heterozygous plasmid pairs (consisting of a wild-type and a mutant plasmid) would correspond to the heterologous plasmid pairs of two different plasmid types. Second, while in some cases, immobilization of plasmid clusters has been observed (Derman *et al*., 2008), others describe a much more chaotic behavior in which plasmids (or plasmid clusters) diffuse in the cell and are over the course of the cell cycle repeatedly repelled by filaments that dynamically form and dissolve (Campbell and Mullins, 2007; Sengupta *et al*., 2010). This would presumably lead to a mixup of plasmid replicates within the cell. Interestingly, based on this observation, Anand and Khan (2010) explicitly point to the “possibility of random assortment of daughter plasmids (sisters and nonsisters) during cell division”. In that case, our assumption of random segregation of wild-type and mutant plasmids would be reasonable.

As a last remark, we would like to stress that a copy number of one (or two, three…) is usually meant to correspond to one plasmid copy per genome equivalent, and often more than one copy is present in the cell even for single copy plasmids. We ignore this in our model in the same way as most models for adaptation on the bacterial chromosome assume that bacteria are strictly haploid.

### Further limitations and extensions

As a caveat, we would like to stress that we ignored plasmid costs for most part of the manuscript. The aim was to disentangle immediate consequences of the plasmid copy number on the dynamics without conflating factors. The burden imposed by plasmids on their bacterial host is expected to increase with the plasmid copy number and at some point outweighs the benefits. Hence, for dominant mutations, an intermediate number of plasmids is most beneficial. The same holds true for adaptation in genes with a gene dosage effect, since the positive effect of a larger number of plasmids in reality saturates (unlike in our model that approximates the fitness benefits of additional plasmid copies by a linear function). However, it is important to note that plasmid costs are often alleviated by compensatory mutations such that they can be very low, undetectable, or even absent (although it is hard to imagine that they could be zero with an ever increasing copy number); see e.g. San Millan *et al*. (2014); Harrison *et al*. (2015); Loftie-Eaton *et al*. (2017).

In our model of evolutionary rescue, we only considered two environments and an abrupt switch between them. Yet, in the context of resistance evolution, it would be highly relevant to include the pharmacokinetics of the drug and to study effects of treatment factors such as the frequency of drug administration on resistance evolution. While the general approach is suitable to deal with more complex scenarios, the mathematical analysis is restricted to constant environments. Considering time-varying drug pressures would require to derive time-dependent establishment probabilities based on time-inhomogeneous branching processes.

Importantly, we assumed that the plasmids are non-transmissible. For transmissible plasmids, horizontal gene transfer could potentially have a non-negligible effect on the establishment of mutated plasmid copies. Conjugative plasmids usually have rather low copy numbers (< 10) since the genes required for conjugation are costly (Norman *et al*., 2009). Transmissible plasmids with higher copy numbers could still co-transfer with other plasmids or phages (Eberhard, 1990; Barry *et al*., 2019). Horizontal transfer is mainly relevant if the population consists of plasmid-carrying and plasmid-free cells since conjugative transfer of plasmids into cells that already contain plasmids of the same incompatibility group (i.e. especially copies of its own type) is unstable or is even prohibited through surface exclusion proteins (Eberhard, 1990).

Last, throughout the article, we assumed that novel beneficial alleles arise by mutation. If instead genes are picked up by transformation and integrated into one of the plasmid copies, the rate of acquisition of beneficial traits is independent of the copy number. Hence, it is only the establishment probability that determines which copy number maximizes the probability of rescue. From our results for the establishment probability, it is straightforward to see that in that case, rescue becomes less likely with increasing copy numbers unless there are gene dosage effects. This result is supported also by simulations done by Ilhan *et al*. (2019) comparing the fixation dynamics of beneficial alleles on multicopy plasmids. In their study, the proportions of populations where the beneficial plasmid allele gets lost by random fluctuations is higher for 100 plasmids per cell compared to 10 plasmids per cell given the total number of mutated plasmids is the same at the beginning (compare the results for p10×10^5^ with those for p100×10^5^ in Fig. 3 in Ilhan *et al*. (2019)).

### Conclusion

Multicopy plasmids are pervasive in bacteria, influencing their ecology and evolution. To capture the population genetics of bacteria, theory therefore needs to account for the contribution of plasmids that are present in the bacterial cell in several copies. Such theory needs to describe the segregation process and take the – possibly environment-dependent – dominance of plasmid-carried alleles into consideration. We here developed and analyzed a model for evolutionary rescue through mutations on multicopy plasmids. Our analysis provides insight into the factors governing the adaptive process and how they balance for various dominance functions and gene dosage effects. Overall, the results demonstrate the relevance of the plasmid copy number for the dynamics of plasmid-carried alleles and for plasmid-mediated bacterial adaptation.

## Acknowledgements

The authors thank Sebastian Bonhoeffer for helpful discussions and support and Alvaro San Millan for providing valuable insights into the biology of multicopy plasmids and Javier Lopez-Garrido for pointing out the existence of negative gene dosage effects. M.S. is a member of the International Max Planck Research School for Evolutionary Biology and gratefully acknowledges the benefits provided by the program.

## A Calculating the establishment probability from branching process theory

To determine the establishment probability of the resistance mutation, we use the mathematical theory of multitype branching processes. A “type” is determined by the plasmid composition of the cell. With a plasmid copy number of *n*, there are hence *n* + 1 types, since cells can contain 0, 1, … *n* mutated plasmids. In the following, we speak of cells of type *i* if a cell contains *i* mutant plasmids. At cell division, a cell produces cells of types that may differ from its own. Only homozygous cells exclusively produce cells of their own types. As a consequence, unless the branching process goes extinct, a permanent lineage of mutant homozygotes will establish. We denote by 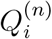 the probability that a cell lineage that is founded by a single cell of type *i* goes extinct. Since the alternative to extinction is establishment of the resistance mutation, it holds

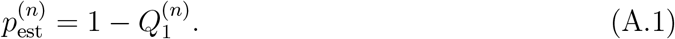

While our model is formulated in continuous time, we consider a discrete-time branching process. This is possible since within our framework with time-independent birth and death rates, it does not matter for establishment or loss of the allele *when* cells divide but only *if* they divide. It is therefore sufficient to focus on whether the next event in the life of a cell is cell division or cell death. The respective probabilities for a cell of type *i* are given by 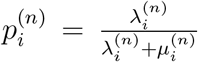 and 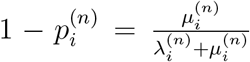. Given the cell divides, it produces cells of type *k* and (2*i + x*) − *k* with probability *P*(*i* → {*k*, (2*i + x*) − *k*}) as defined in Eq. (1). Transitions between types in the branching process therefore occur with probabilities 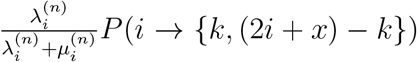. Since the mutation probability *u* is small, we can neglect new mutations during the establishment process. We therefore set *x* = 0 in the branching process approximation.

Following these considerations, it is intuitive to derive a set of (coupled) equations for the probabilities 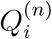. The lineage founded by a single cell of type *i* goes extinct either if the cell dies or if the lineages founded by its two daughter cells both go extinct. Thus:

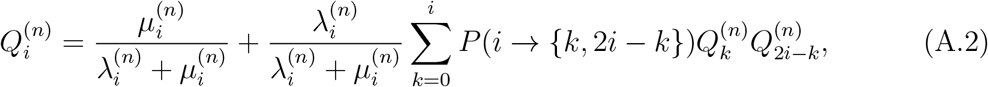

Note that since we ignore new mutations during the establishment process, we have 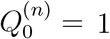,

In the following, we provide a brief formal summary of how to determine the vector 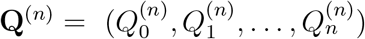. This follows standard theory on multitype branching processes, and we refer to textbooks for details (Mode (1971), Haccou *et al*. (2005)).

We denote the probability generating function of the multitype branching process by

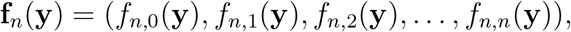

where the vector components *f_n,i_*(**y**) are given by

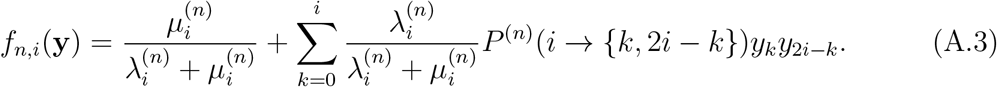

The vector **y** = (*y*_0_, *y*_1_, …, *y_n_*) is a dummy variable without biological meaning. The probability generating function can be used in two different ways to determine 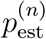. In our model analysis below, we will make use of both methods.

First, the vector **Q**^(*n*)^ of extinction probabilities is a fixpoint of the probability generating function (Mode (1971)), i.e.

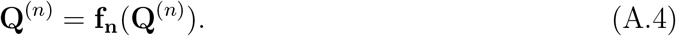

This exactly corresponds to the set of equations (A.2). We use the fixpoint equation to obtain the extinction probabilities for *n* = 1 and *n* = 2.

Second, it holds that (Eq. (7.4) in Mode (1971))

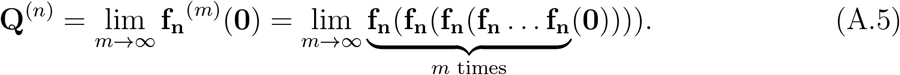

We apply this to numerically obtain the extinction probabilities for *n* > 2.

From Eq. (A.2), one immediately obtains for the establishment probability of a process that is initiated by a single mutant homozygote:

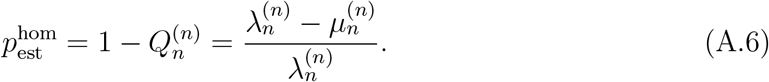

## B Model linking the dominance function to the antibiotic concentration

In this section, we provide a detailed presentation of the model that connects the fitness of a specific cell type to the antibiotic concentration in the environment and that underlies Fig. 3 in the main text.

For the bacterial response to a given antibiotic concentration *within the cell, c*_in_, we use a pharmacodynamic function (cf. Regoes *et al*. (2004)). We assume that the net growth of a bacterial population in dependence of the internal antibiotic concentration is given by

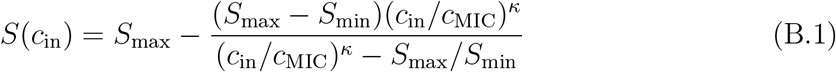

The four parameters denote maximal and minimal net growth (*S*_max_ and *S*_min_), the antibiotic concentration level *c*_MIC_ where the net growth is zero and the Hill coefficient *κ*, which determines the slope of the curve. As we will see below, the concentration within a cell depends on its type, i.e. 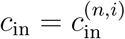. We set the death rate 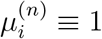 for all *n* and *i* and the birth rate as 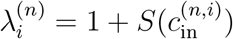.

The rate of antibiotic inflow *κ*_in_ into the cell is proportional to the difference in the concentrations outside and inside the cell *c*_ou_t and *c*_in_, i.e.

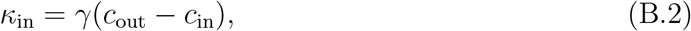

where *γ* denotes the diffusion coefficient.

Within the cell, the antibiotic can be degraded by enzymes that are encoded by the mutant plasmid. The degradation rate *κ*_deg_ depends on the antibiotic concentration in the cell and the amount of degrading enzymes that can break antibiotic molecules and that depends on *n* and *i*,

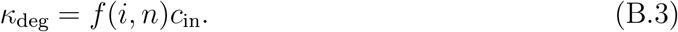

In the main text, we consider two cases. In one case, the amount of available enzymes is proportional to the relative number of mutant plasmids,

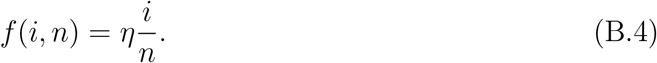

In the other case, it is proportional to the absolute number of mutant plasmids,

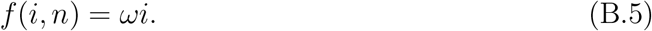

We assume that inflow of antibiotic molecules into the cell and degradation of antibiotics are in equilibrium:

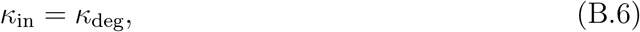

From this, we obtain the concentration 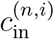 within cells that carry *i* mutant and *n − i* wild-type plasmids, given an external concentration *c*_out_:

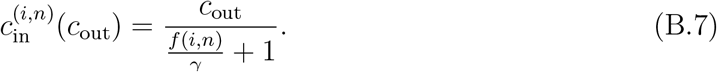

Substituting equation (B.7) in (B.1), we obtain the net growth 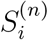 as a function of the plasmid composition.

## C Modeling the cell type frequencies in the standing genetic variation

To obtain the cell type frequencies in the standing genetic variation we set up a multitype Moran that describes a bacterial population of constant size *N*_0_ prior to antibiotic treatment. When a cell divides, the daughter cells replace the parental cell as well as another randomly chosen cell from the population. Wild-type cells replicate at rate 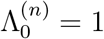. Since resistance mutations may cause a cost in the antibiotic-free environment, we assume that replication of cells carrying mutated plasmids occurs at a lower rate. We set for *i* > 0:

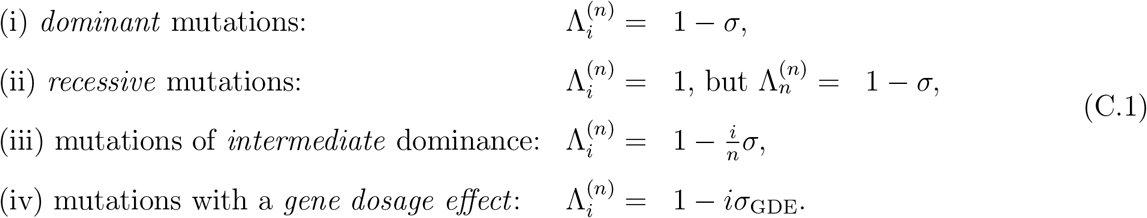

Mutation, plasmid replication, and plasmid segregation happen in the same way as described in the main text for the phase after the environmental change.

For the stochastic simulations, the model is implemented in a straightforward way. The initial population consists of *N* wild-type cells. Transitions between states may happen when a cell of type *i* divides, another cell of type *k* is replaced and two daughter cells enter the population. We denote the types of the first daughter cell by *j*_1_ and of the second daughter cell by *j*_2_ (note that the cells are ordered now unlike in Eq. (1)). The reaction can then be expressed by

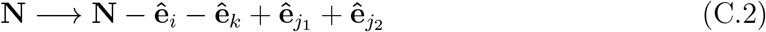

The rate of this reaction is given by

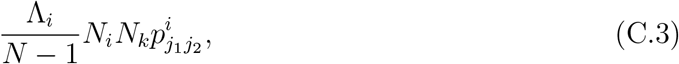

where 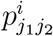 denotes the probability that the first daughter cell of a type *i* cell is of type *j*_1_ and the second one of type *j*_2_:

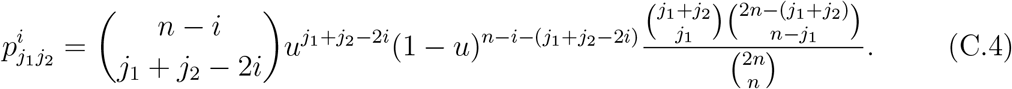

We simulate this process, using the Gillespie algorithm (Gillespie, 1976) which we implemented in the Python programming language. At simulation time *t*_sim_ = 1000, the switch in the environment occurs and we continue the simulations according to the model dynamics in the stressful environment described in the main text to determine the probability of evolutionary rescue. In order to obtain the frequencies of the various cell types in the standing genetic variation, we record those at time *t*_sim_ = 1000.

We next derive a set of ordinary differential equations that describes the dynamics of the systems deterministically. To proceed, we look at all the events at which the number of cells of type (*n, i*) changes. We first omit all new plasmid mutations.

- Case I: a cell of type *k ≠ i* reproduces

**– A:** one of the daughter cells is of type *i*; then:

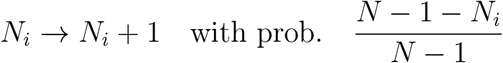
**– B:** neither of the daughter cells is of type *i*; then

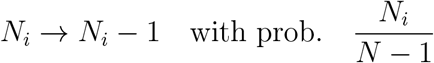 Note that it is not possible that both daughter cells are of type *i* since there are 2*k* mutant plasmids before cell division, and *k ≠ i* for the case considered here.
- Case II: a cell of type *i* reproduces

**– A:** both daughter cells are of type *i*; then

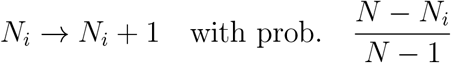
**– B:** neither of the daughter cells is of type *i*; then

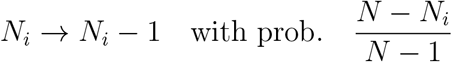

and

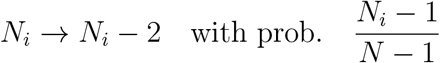 Note that for similar reasons as above, it is not possible that a cell of type *i* has only one daughter cell of type *i* and one of another type.

Putting everything together, we obtain for the dynamics of cells of type *i* (where we use the shorthand notation 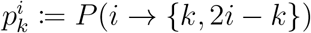, see Eq. 1):

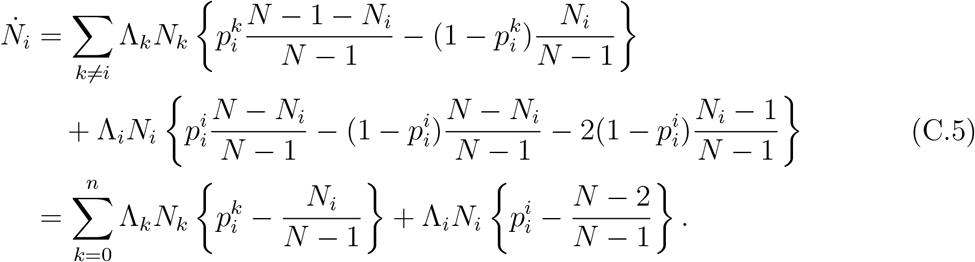

We now modify Eq. (C.5) to include new mutations of wild-type to mutant plasmids. As in the approximation for the establishment process, we only consider mutations at replication of wild-type cells with *i* = 0. For other cells, the dynamics of mutant plasmids are governed by segregation and new mutations can be neglected. Furthermore, within one cell division it is highly unlikely that more than one mutation occurs. Consequently, only the dynamics of cells with *i* = 0 or *i* = 1 mutated plasmids are affected by mutations. We approximate the probability that no mutation occurs by 1 − *un*, and the probability that one mutation occurs by *un*.

For *i* = 0, case II (reproduction of cell of type 0) from above is altered:

- **A**: both daughter cells are of type 0 (i.e. no mutation); then

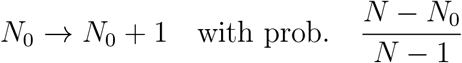

this happens with probability (1 − *un*)
- **B**: daughter cells are of type 0 and 1 respectively (one mutation); then

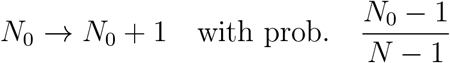

this happens with probability *un*

Therefore, including plasmid mutations, Eq. (C.5) for *i* = 0 gets

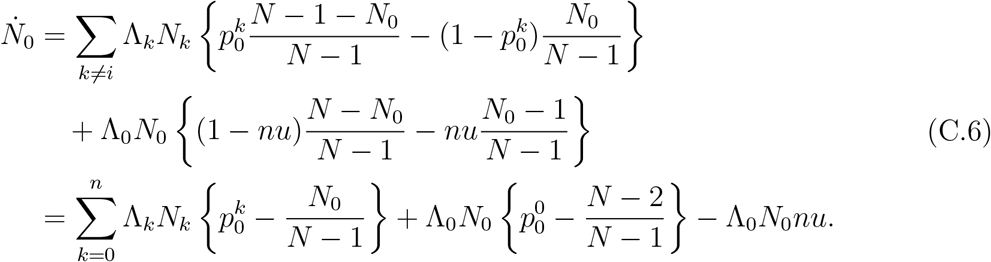

Comparing Eqs. (C.5) and (C.6), shows that plasmid mutations lead to a decrease in frequency of wild-type cells (*i* = 0) with a rate λ_0_*N*_0_*nu*. This will increase the frequency of cells of type (*i* = 1). Hence, we modify the dynamics of *N*_1_ with the same but positive rate:

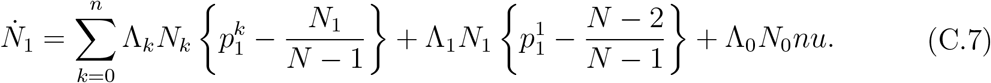

We obtain the frequencies *x_i_* of all cell types *i* ∈ {0, …, *n*} in the standing genetic variation of an infinite population *N* → ∞ by numerically integrating the non-linear system 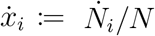 with 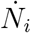 given by Eq. (C.5), (C.6), and (C.7) starting with *x*_0_ = 1, *x*_i_ = 0 for *i* > 0. For this purpose, we use the `solve_ivp` method from the SciPy package in Python. The equilibrated frequencies 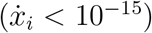 are used to estimate the average number of cell type abundances *N_i_ = N_x_i__* for a given population size *N*.

