## Supplementary Information for "Evolutionary rescue and drug resistance on multicopy plasmids"

### Supporting information – Evolutionary rescue and drug resistance on multicopy plasmids

#### S1: Alternative models of plasmid replication and segregation

In this section, we present two alternative models of plasmid replication and segregation. For the model described in the main text, we assume that prior to cell division each plasmid is replicated once and all copies are randomly distributed to the daughter cells, where each daughter cell reveives $n/2$ copies. For the alternative models, we assume that at cell division plasmids are distributed to both daughter cells *before* they replicate to reach their copy number. We consider two variants: (1) each cell receives $n/2$ plasmid copies for $n$ even or $(n-1)/2$ and $(n+1)/2$ if $n$ is uneven (2) each daughter cell receives at least one plasmid; the remaining plasmids are randomly distributed to the daughter cells such that in general, daughter cells receive different numbers of plasmid copies. Subsequently, plasmids get replicated one plasmid at a time until there are $n$ plasmid copies in the cell. The plasmid copy that is replicated is chosen randomly from all plasmids in the daughter cell (including those that have just been generated). Therefore, the plasmid composition changes during the plasmid replication phase. Moreover, an early mutation appearing during the replication phase can immediately lead to a daughter cell with a number of mutated plasmids greater than $1$.

Fig. S1.1 shows the probabilities of *de novo* rescue under the two alternative models for the same parameter combinations as Fig. 2. The results obtained from the two variants of the alternative model are very similar to each other. In Fig. S1.2, we directly compare the results from the alternative models to the original model from the main text. While the general trends mostly remain the same, some differences arise. The most prominant deviations arise for recessive mutations. Under the original model, the rescue probability drops very quickly with increasing plasmid copy number. With the alternative schemes of plasmid replication and segregation, a high copy number has a less negative effect on rescue, and there is even a slight maximum for low copy numbers. Likewise, for mutations of intermediate dominance, the negative effect of high copy numbers is also weaker. In that case, this makes that we do not observe the slight intermediate maximum anymore that we observed under the original model.

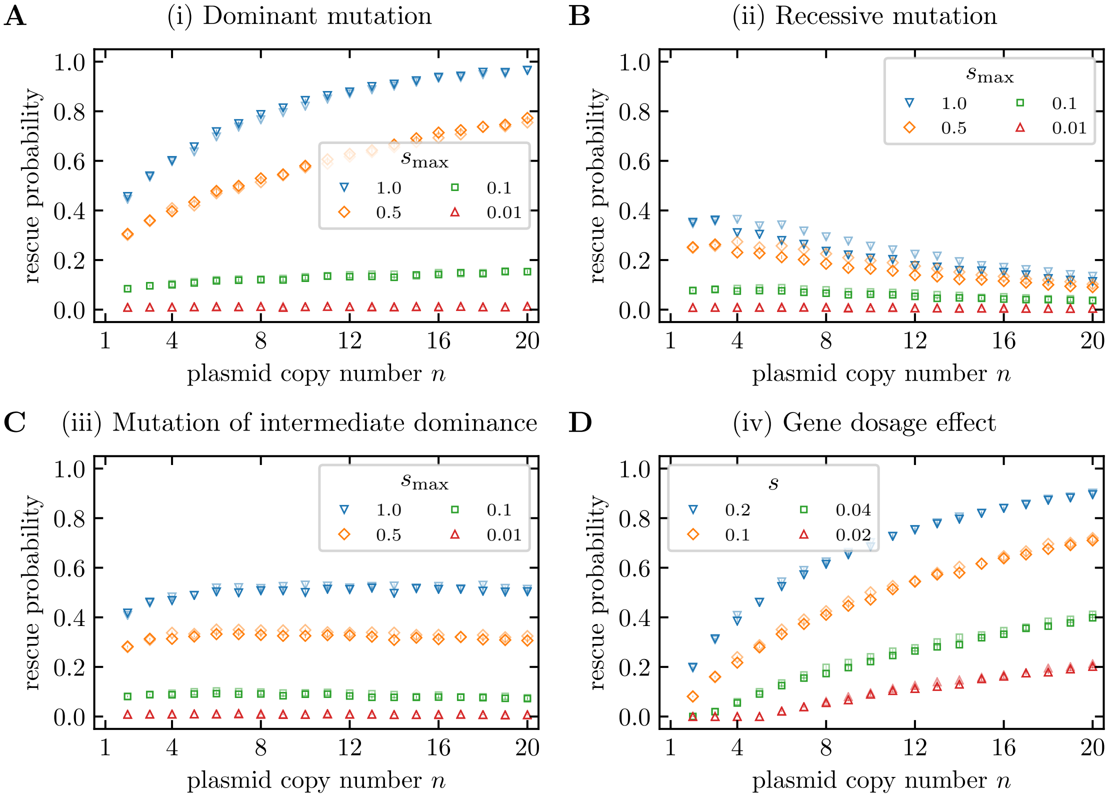


**Figure S1.1**: **Rescue probabilities under two alternative schemes of plasmid replication and segregation.** For both schemes, plasmids are at first segregated into the daughter cells, and subsequently plasmids get replicated one by one until the cell contains $n$ plasmid copies. For alternative 1 (dark colored markers), plasmid are segregated to the daughter cells in equal numbers $n/2$ if $n$ is even and one daughter cell receives one more plasmid than the other if $n$ is uneven. For alternative 2 (light colored markers), each daughter cell receives at least one plasmid copy, and the remaining copies are segregated randomly. The parameters correspond to those in Fig. 2A-D in the main text. Results were obtained by $10^4$ stochastic simulations ($10^5$ for alternative 1 and $n \leq 10$). Error bars indicate standard errors.

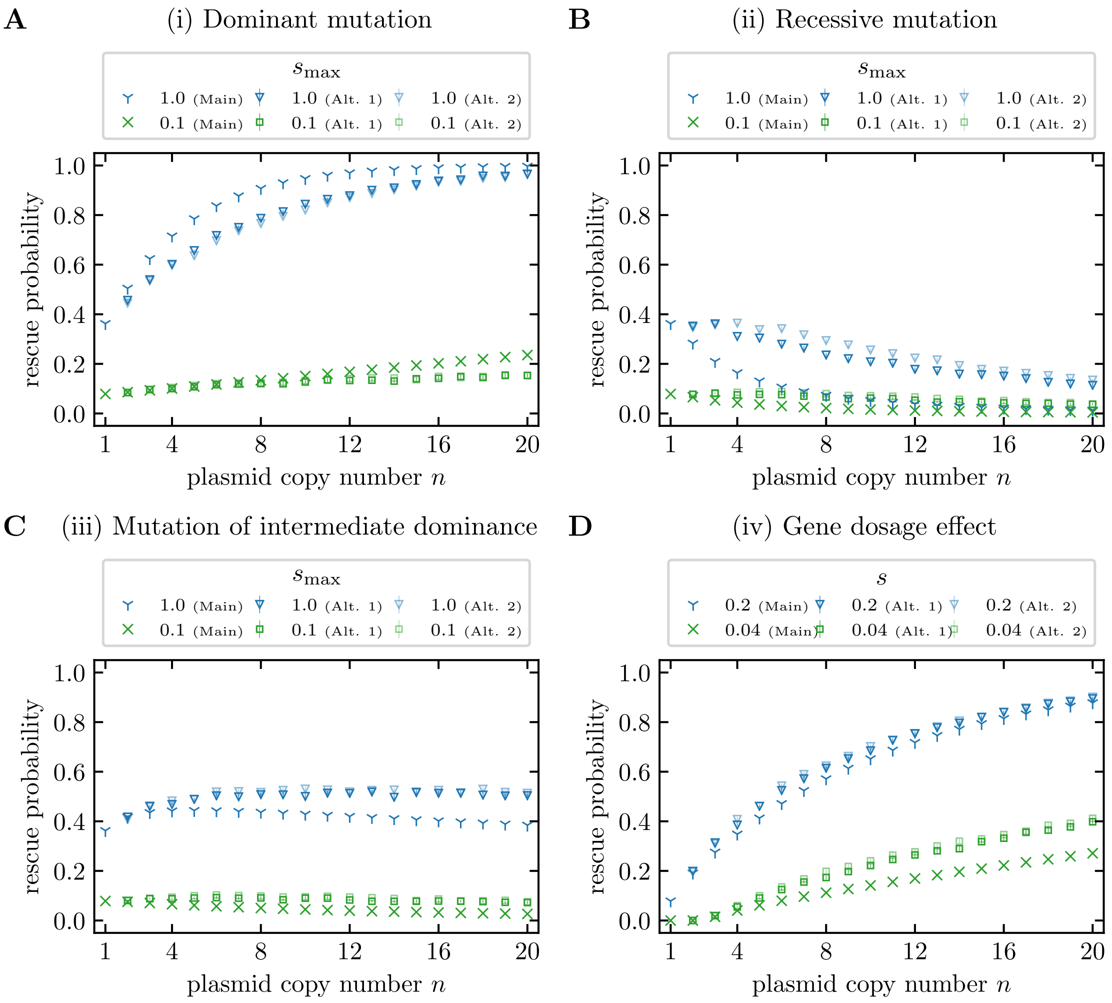


**Figure S1.2**: **Comparison of rescue probabilities from the main text model with results obtained from the alternative models.** The alternative models are described in the text and in the caption of Fig. S1.1. As a reminder, for alternative 1 (dark open colored markers), plasmid are segregated to the daughter cells in equal numbers $n/2$ if $n$ is even and one daughter cell receives one more plasmid than the other if $n$ is uneven. For alternative 2 (light open colored markers), each daughter cell receives at least one plasmid copy, and the remaining copies are segregated randomly. The parameters correspond to those in Fig. 2A-D in the main text and in Fig. S1.1 in the SI. Results for the main text model were obtained numerically by using Eq. (3) with Eq. (A.5) and Eq. (A.1). For the alternative models, results were obtained by $10^4$ stochastic simulations ($10^5$ for alternative 1 and $n \leq 10$). Error bars indicate standard errors.

### S2: Comparing rescue with one and two plasmid copies and gene dosage effects

In this section, we provide the mathematical proof for the statement (made in the maintext) that the probability of evolutionary rescue $P_{\text{rescue}}$ is higher for $n = 2$ plasmids per cell compared to $n = 1$ in a scenario with a gene dosage effect (scenario (2)). As a reminder, in this scenario the level of resistance, given by birth rates $\lambda_i^{(n)}$ and death rates $\mu_i^{(n)}$, depends on the number of mutated plasmids $i$. Therefore we use birth rates $\lambda_0$, $\lambda_1$, $\lambda_2$ and death rates $\mu_0$, $\mu_1$, $\mu_2$ independently of the plasmid copy number $n$ as defined in the main text. Furthermore, we define the ratio of death and birth rates $\rho_1 \equiv \frac{\mu_1}{\lambda_1}$ and $\rho_2 \equiv \frac{\mu_2}{\lambda_2}$.

It needs to be shown that the rescue probability given by Eq. (3) for $n = 2$ is higher than for $n = 1$. This is equivalent to

$$1 - e^{-2u\lambda_0 \frac{N_0}{\mu_0 - \lambda_0} p_{\text{est}}^{(2)}} \overset{!}{>} 1 - e^{-u\lambda_0 \frac{N_0}{\mu_0 - \lambda_0} p_{\text{est}}^{(1)}}$$

which can be transformed into the following inequality:

$$2p_{\text{est}}^{(2)} \overset{!}{>} p_{\text{est}}^{(1)}. \tag{S2.1}$$

In the 'worst' case, the second plasmid brings no additional benefit, i.e. $\rho_2 = \rho_1 \equiv \rho < 1$. In this scenario, establishment probabilities for one and two plasmids per cell (see Eq. (4)) become

$$\begin{aligned} p_{\text{est}}^{(n=1)} &= 1 - \rho, \\ p_{\text{est}}^{(n=2)} &= \frac{1}{4} - \frac{3}{4}\rho + \frac{1}{4}\sqrt{9 - 14\rho^2 + 9\rho^2}. \end{aligned}$$

Inserting the establishment probabilities in Eq. (S2.1) shows that the statement is correct:

$$\begin{aligned} & \frac{1}{2} - \frac{3}{2}\rho + \frac{1}{2}\sqrt{9 - 14\rho^2 + 9\rho^2} > 1 - \rho \\ \Rightarrow\ & \sqrt{9 - 14\rho^2 + 9\rho^2} > 1 + \rho \\ \Rightarrow\ & 9 - 14\rho^2 + 9\rho^2 > 1 + 2\rho + \rho^2 \\ \Rightarrow\ & 1 - 2\rho + \rho^2 > 0 \\ \Rightarrow\ & (1 - \rho)^2 > 0. \end{aligned}$$

### S3: The effect of the variance in the cell type numbers in the standing genetic variation

Fig. S3.3 shows the probability of rescue from the standing genetic variation with all parameters being equal as in Fig. 5 except for that the population size is smaller by an order of magnitude. Deviations between the analytical theory and the stochastic simulations are larger for smaller populations.

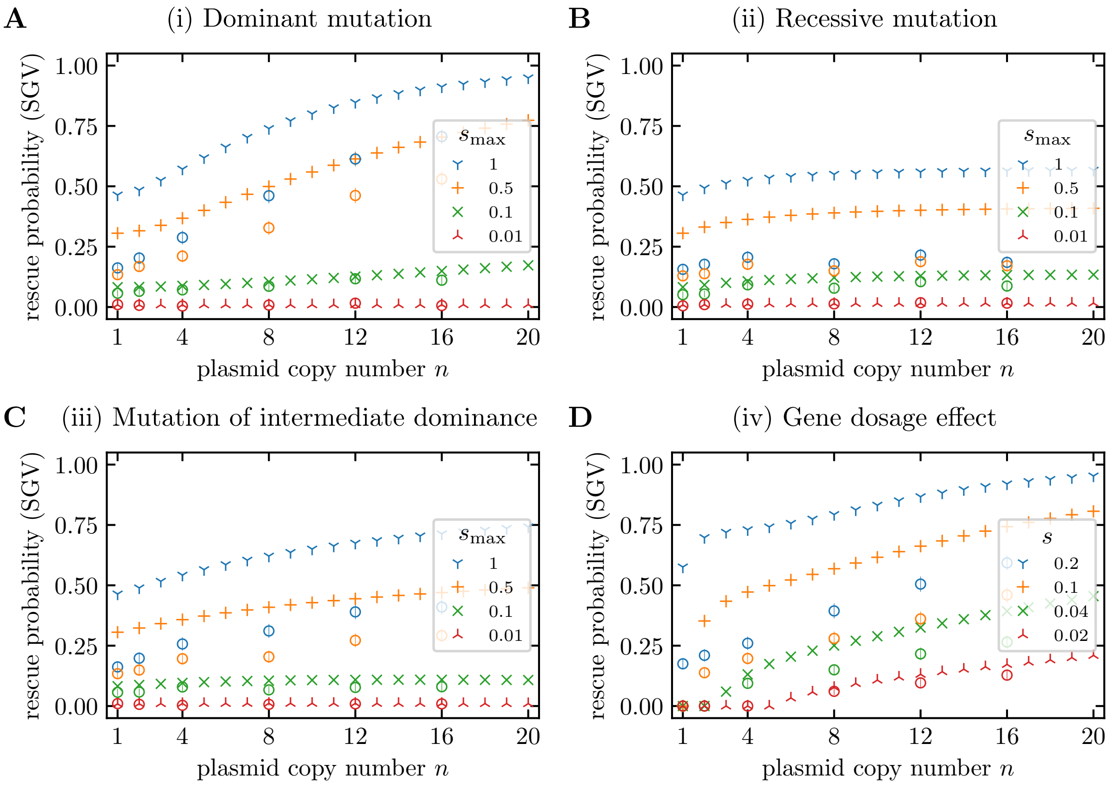


**Figure S3.3**: **Probabilities of evolutionary rescue from standing genetic variation.** The figure corresponds to Fig. 5 in the main text except for a smaller population size of $N_0 = 3 \times 10^8$ compared to $N = 3 \times 10^9$ in Fig. 5.

In the main text, we hypothesized that the decrease in $P_{\text{rescue}}^{(\text{SGV})}$ for large $n$ for recessive mutations is caused by a higher variance in the number of cells $N_i$ with high $n$. Here, we investigate this in a bit more detail. To this purpose, we choose to consider a scenario where the wildtype (and hence all heterozygous cells) are lethal in the new environment. Hence, rescue – if it occurs – occurs from mutant homozygotes in the standing genetic variation. Fig. S3.5A shows the frequency of mutant homozygotes in dependence of the plasmid copy number, and Panel B shows their variance. One can see that the mean frequency

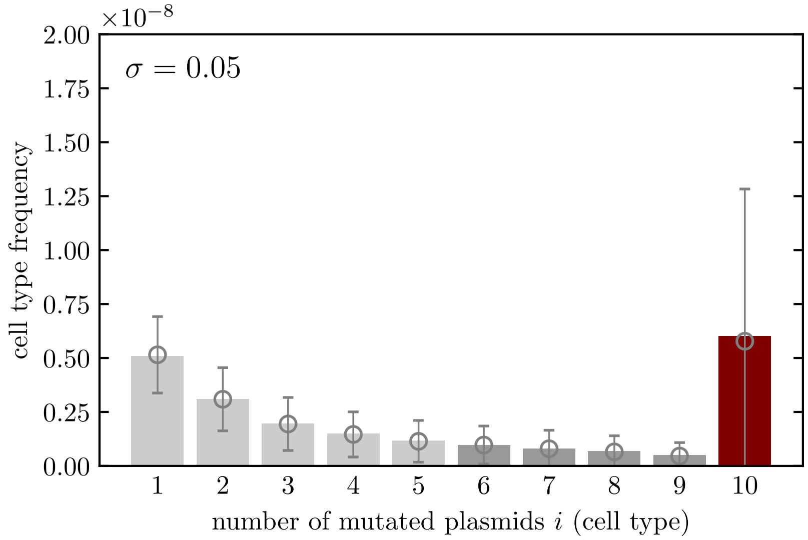


**Figure S3.4**: **Cell type frequencies in the standing genetic variation.** The figure reproduces Fig. 4F with a larger range of the y-axis such that the entire error bar for the frequency of the homozygous type (red bar) is visible.

remains constant but the variance indeed increases with $n$. In line with $N_n = \text{const.}$, the analytical theory predicts that $P_{\text{rescue}}^{(\text{SGV})}$ does not change with $n$ (Fig. S3.5). In contrast, the results obtained from stochastic computer simulations decline as $n$ increases, which confirms our intuition.

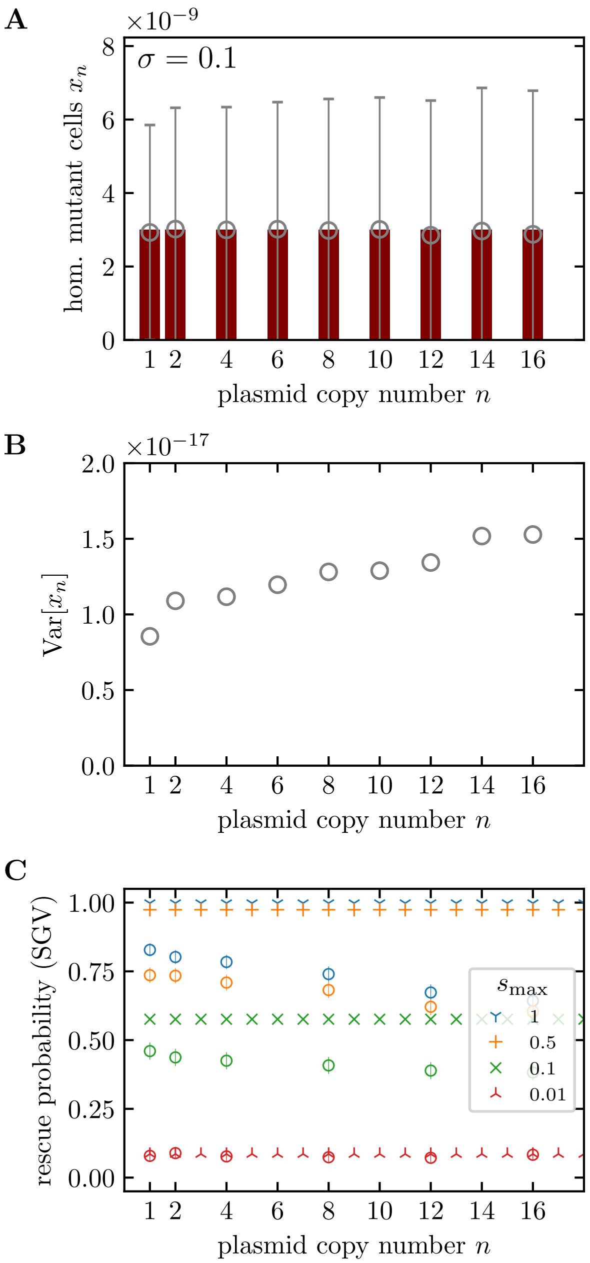


**Figure S3.5**: **Influence of stochasticity in the cell type frequencies in the standing genetic variation on the probability of evolutionary rescue.** Panel A: Frequencies of homozygous mutant cells $x_n = N_n/N$ for various plasmid copy numbers obtained from the deterministic mutation-selection equilibrium (bars) and from stochastic simulations (circles) for a recessive mutation with selection coefficient $-\sigma$. Error bars indicate the standard deviation $\sigma_{x_n} = \sqrt{\mathrm{Var}[x_n]}$ (*not* the standard error!). Panel B: Variance $\mathrm{Var}[x_n] = \sigma^2_{x_n}$ of the number of homozygous mutant cells $x_n$ (cf. error bars in Panel B). Panel C: Probability of evolutionary rescue from standing genetic variation considering a scenario where heterozygous cells (as well as wild-type cells) are lethal. All parameters are the same as in Fig. 5 (main text) except for the birth rate of wild-type and heterozygous cells $\lambda_0^{(n)} = 0$ in the presence of antibiotics. Simulation results are obtained from $10^3$ replicate simulations.
